# Generation of functional vasculature from engraftable human pluripotent stem cell-derived progenitors

**DOI:** 10.64898/2026.05.14.723516

**Authors:** Ian M. Fernandes, Hao Yin, Yuan Yao, Blair K. Gage, Zengxuan Nong, Mark Gagliardi, Molly Shoichet, Geoffrey Pickering, Gordon Keller

**Affiliations:** McEwen Stem Cell Institute, University Health Network, Toronto, ON M5G1L7, Canada; Department of Medical Biophysics, University of Toronto, Toronto, ON M5G1L7, Canada; Princess Margaret Cancer Center, University Health Network, Toronto, ON M5G1L7, Canada; Department of Chemical Engineering and Applied Chemistry, University of Toronto, Toronto, ON M5S 3E5, Canada; Donnelly Centre for Cellular and Biomolecular Research, University of Toronto, Toronto, ON M5S 3E1, Canada; Institute of Biomedical Engineering, University of Toronto, Toronto, ON M5S 3G9, Canada; Robarts Research Institute, Schulich School of Medicine and Dentistry, Western University, London, Canada; Department of Medical Biophysics, Schulich School of Medicine and Dentistry, Western University, London, Canada; Department of Biochemistry, Schulich School of Medicine and Dentistry, Western University, London, Canada; Department of Medicine, Schulich School of Medicine and Dentistry, Western University, London, Canada; London Health Sciences Centre London, Canada; Sprott Centre for Stem Cell Research, Ottawa Hospital Research Institute, Regenerative Medicine Program, Ottawa, ON K1H8L6, Canada

## Abstract

The ability to revascularize target tissues and organs through cell-based therapy would provide a novel approach for the treatment of a range of ischemic disorders including cardiovascular diseases, stroke and peripheral artery disease. Towards this goal, we have identified a human pluripotent stem cell (hPSC)-derived vascular progenitor (VP) population generated via an epicardial intermediate with functional engraftment properties. VP cells efficiently engraft the mammary fat pad and hind limb skeletal muscle of NSG recipient mice and form vessel-like structures that integrate with the host vasculature. In an ischemic hind limb mouse model, VPs generate extensive vascular grafts that improve perfusion, restore some function and preserve muscle integrity over a three-month period post-transplant. Single-cell transcriptomic and flow cytometric analyses show that the VP population, initially identified by the co-expression of CD140b, CD13 and KDR, displays an epicardial lineage signature and expresses a spectrum of genes and proteins indicative of vascular progenitor stage cells. Together, these findings demonstrate that it is possible to revascularize both normal and ischemic tissue through the transplantation of an appropriate hPSC-derived progenitor and in doing so, lay the foundation for developing cell-based therapy approaches to treat ischemic diseases.

**Graphical Abstract Legend:** Human pluripotent stem cells are differentiated through an epicardial intermediate to generate vascular progenitor (VP) cells characterized by expression of CD140b, CD13 and KDR. These VP cells demonstrate the capacity to engraft both mammary fat pad and skeletal muscle tissue where they form stable perfused vascular networks. In a hindlimb ischemia model, VP cell transplantation restores blood flow and improves functional outcomes.

**eTOC Blurb:** Fernandes et al. develop a protocol to generate engraftable vascular progenitors from human pluripotent stem cells through an epicardial intermediate. These cells form functional vessels *in vivo*, restore perfusion in ischemic tissue, and demonstrate tissue-specific adaptation while maintaining endothelial identity, providing a foundation for therapeutic revascularization.

**Highlights:**

- A staged differentiation protocol generates vascular progenitors (VPs) from hPSCs via an epicardial intermediate.
- VP cells form stable, perfused vascular networks following transplantation into multiple tissue sites.
- VP cell therapy with or without VEGF nanoparticles restores perfusion and improves functional outcomes in hindlimb ischemia.
- Single-cell analysis reveals tissue-specific adaptation while maintaining endothelial identity.

## Introduction

The blood vascular system, made up of a network of veins, arteries and capillaries, functions to transport blood cells that deliver oxygen and nutrients to all tissues in the body (Augustin & Koh 2024). The vessels of the system, the veins and arteries are comprised of vascular cells including an inner lining of vascular endothelial cells surrounded by a layer of mural cells (pericytes and smooth muscle cells) that provide structural support and regulate vascular tone, blood flow, and vessel stability (Armulik et al., 2005; Armulik, et al.,2011). Diseases that impair vascular function can restrict blood flow, resulting in an ischemic state that can severely compromise organ function. Ischemia represents one of our greatest clinical challenges, as it is the underlying cause of most cardiovascular diseases and stroke, leading causes of death worldwide (Martin et al., 2025). Compromised blood flow also impacts the function of other tissues, including the skeletal muscle of the peripheral limbs leading to a condition known as peripheral arterial disease (PAD). Patients with PAD have severely restricted circulation to the extremities resulting in extreme pain, poor wound healing and in advanced cases, limb amputation (Golledge J. 2022; Norgren et al. 2007; Annex BH 2013). PAD is not rare as it is estimated that more than 200 million people suffer from this condition worldwide (Fowkes et al. 2013; Norgren et al. 2007). The current standard of care for PAD includes medication for pain management and prevention of disease progression and in severe cases, surgical revascularization. While these therapeutic interventions can be effective, they do not provide long-term benefit for many patients and, as a consequence there is an urgent unmet need for new therapies.

Cell-based therapy designed to generate new functional vasculature from transplanted progenitor cells would represent a novel and potentially curative therapy for PAD and other ischemic diseases (Cooke & Losordo 2015; Qadura et al. 2018; Katagiri et al. 2020). The challenge with this approach, however, has been the identification of an appropriate progenitor/vascular cell population that is able to efficiently engraft ischemic muscle tissue and form sufficient new vasculature to restore blood flow and function to the injured tissue. Numerous studies over the past two decades have described different human vascular and mesenchymal cell types that display some capacity to restore vascular function when transplanted in pre-clinical mouse models of hindlimb ischemia (Tateishi-Yuyama et al., 2002, Qadura et al. 2018; Cooke & Losordo 2015; Benoit et al. 2013). In most cases, these populations were isolated from adult tissue, including the bone marrow (Tateishi-Yuyama et al., 2002); adipose tissue (Planat-Benard et al. 2004; Mazo et al. 2008; Ii et al. 2005); and peripheral blood (Asahara et al. 1997). While engraftment was observed early following transplantation, contribution to new vessel formation and vascular development over the long-term was exceptionally low or not detected. Despite limited engraftment, the transplanted cells did provide benefit, including improved perfusion, decreased fibrosis and in some cases, salvaged limbs. Restoration of function in the absence of extensive donor-derived vascularization strongly suggests these effects are not due to new vasculature but rather mediated through a paracrine mechanism, with the transplanted cells transiently secreting factors or stimulating the secretion of factors that promote angiogenesis and/or improve endogenous vascular function. While paracrine-based strategies provide one avenue for developing new regenerative therapies, their ability to sustain long-term benefit has not been well documented. Additionally, this approach will likely have limited efficacy in patients whose own vasculature is compromised due to disease. In such patients, new vasculature generated from transplanted healthy progenitors would conceptually be a more effective therapy.

The challenges faced with revascularizing muscle tissue with adult-derived progenitors/vascular cells likely reflect the fact that these cells are not programmed to generate new vasculature *de novo*. In the adult, new vasculature is primarily generated through a process known as angiogenesis, that involves sprouting of new vessels from existing vasculature (Carmeliet P 2003; Augustin & Koh 2024). Angiogenesis is mediated by mature endothelial cells, not progenitors. By contrast, the vasculature that forms in the tissues and organs during embryonic/fetal life is generated from endothelial and smooth muscle cells derived from immature progenitors via a developmental process known as vasculogenesis. These vascular progenitors, specified from different subsets of mesoderm rapidly differentiated to endothelial progeny that migrate into the developing tissue where, in response to environmental cues give rise to the specified cell types that make up the arterial, venous and capillary vasculature (Trimm & Red-Horse, 2023; Bovill & Spencer, 2023; Barnett et al., 2024; Lin et al., 2026).

The remarkable potential of embryonic/fetal vascular progenitors positions them as ideal candidates for new revascularization cell-based therapies. Pursuing this strategy has, however, been challenging, given the limited access to early human embryonic and fetal tissue. Human pluripotent stem cells (hPSCs) represent an alternative and attractive source of such progenitors, as their differentiated progeny provide access to unlimited numbers of vascular cells at different developmental stages (Lin et al., 2017; Cheung et al., 2012; Wang et al., 2020; Colunga et al., 2019; Lee et al., 2024; Gong et al., 2025). Studies over the past decade have described differentiation protocols that promote the generation of different subtypes of vascular cells from hPSCs including arterial and venous endothelial cells and smooth muscle cells (Patsch et al., 2015; Ditadi et al., 2015; Zhang et al 2017; Rosa et al 2019; Gage et al., 2020; Shen et al., 2021). Transplantation studies have shown that hPSC-derived vascular cells can engraft the heart and liver of recipient animals, demonstrating some capacity to function in adult tissue (Fernandes et al 2011; Fernandes et al 2015; Gage et al., 2020; Gage et al., 2022; Wang et al., 2020). Two recent reports have shown that hPSC-derived vascular cells are also able engraft ischemic hind limb muscle in immunocompromised mice and provide some benefit to the injured tissue. In the first of these, Gong et al., transplanted hPSC-derived 3D vascular organoids into ischemic muscle and documented improved perfusion and reduced fibrosis over a 2-week period post-transplant (Gong et al. 2025). Histological analyses revealed the presence of some human vasculature at this early stage. The second study involved the transplantation of single cell suspensions of hPSC-vascular cells into ischemic muscle (Huslein et al., 2026). These animals also showed improved vascular function in the injured tissue which persisted for the 2-month duration of the study. However, while improvements were detected, overall engraftment levels were low and declined with time, strongly suggesting that these effects were largely paracrine mediated. Collectively, this work indicates that hPSC-derived vascular cells can engraft ischemic muscle tissue at least for short periods of time and that this engraftment leads to some improvement in blood flow and tissue damage. Whether or not long-term stable vascular engraftment can be achieved in muscle tissue remains to be determined.

In this study we sought to identify a hPSC-derived transplantable progenitor that would fulfill two key criteria essential for an effective therapy for PAD. The first, is the capacity to engraft target tissue efficiently when delivered as a single cell suspension. The second, is the ability to generate readily measurable human vascular grafts that would integrate with the host vasculature and persist for greater than two months. Here we report on the identification of a hPSC-derived epicardial-derived vascular progenitor (VP) population that fulfills these criteria. These progenitors display the potential to engraft both fat pad and skeletal muscle tissue when transplanted as a single cell suspension and form extensive human vascular grafts that integrate with the mouse vascular system. We also demonstrate that the VPs are able generate vascular grafts in ischemic hind limb muscle for up to 90 days and that these grafts restore blood perfusion and hind limb function and preserve muscle architecture over this period of time. Finally, our transcriptomic and cell surface marker analyses revealed that the VP population is highly enriched in progenitor stage cells and contains few, if any mature endothelial cells. Taken together, these findings show it is possible to revascularize ischemic tissue and that success is dependent on identification of the correct progenitor population.

## Results

### Generation of Epicardial-derived Vascular Progenitor Cells from Human Pluripotent Stem Cells

To generate a vascular progenitor population for transplantation, we modified our previously described protocol designed for the production of epicardial-derived cardiac fibroblasts (Figure 1A) (Fernandes et. al., 2023). This 23-day protocol transitions through distinct developmental stages that include the generation of a pro-epicardial population at day 8 of differentiation (Stage 1), the specification of an epicardial cell population by day 17 (Stage 2) and the production of the desired epicardial derivatives by day 23 (Stage 3). The final step (days 17 to 23) involves an epithelial-to-mesenchymal transition (EMT, days 17-20) followed by a differentiation and expansion step (days 20-23) and depending upon the cytokines used will give rise to either fibroblasts and/or vascular cells with a yield of approximately 3 to 5 cells (+/-1.6 cells) of the target cell type per input epicardial cell. To generate vascular lineage progenitors, we focussed on modifying the factors during the EMT and differentiation/expansion steps reasoning that these are the critical stages for specifying different fates. The same factors used to generate the Stage 1 and Stage 2 populations in the fibroblast protocol were maintained in the current iterations. The resulting day 23 populations were analyzed for the expression of surface markers indicative of vascular progenitor development including the combination of CD140b/CD13 for the perivascular lineage and KDR/CD31 for the endothelial lineage. With this approach, we found that treatment of the day 17 population with the combination of bFGF, PDGF-BB, TGFβ1 and EGF for three days followed the combination of bFGF, PDGF-BB and the TGFβ inhibitor SB431542 (SB) for an additional three days promoted the development of a population that expressed both perivascular and endothelial markers. The inclusion of PDGF-BB was based on its established role in mural cell specification during vascular development (Hellström et al., 1999; Mellgren et al., 2008), while the combination of bFGF, TGFβ1, and EGF was selected to promote EMT based on their known activities in epicardial development (Morabito et al., 2001; Austin et al., 2008; Kirschner et al., 2011). These factor combinations differ from those established for our fibroblast protocol that included retinoic acid and CHIR99021 (WNT Agonist) from days 17-20 and the combination of bFGF, and SB431542 (SB) from days 20-23. As shown in Figure 1B the majority of the cells in the day 23 vascular-induced population expressed CD140b and of these approximately 75% co-expressed CD13. More than half of the CD140b^+^CD13^+^ population also expressed KDR. This CD140b^+^CD13^+^KDR^+^ fraction typically represents greater than 60% of the total population at this stage (Figure 1C). CD31^+^ cells were not detected at this stage. Analyses of the earlier stage populations showed that the majority of cells at both time points expressed CD140b (Fig. 1C). The proportion that co-expressed CD13 increased over this time. KDR showed a dynamic expression pattern, with most cells being positive at stage 1 and fewer than 25% of the day 17 epicardial population retaining expression. Less than 20% of the total stage 1 and 2 populations co-expressed these three markers. A portion of the day 23 fibroblast-induced population also expressed CD140b (Fig 1D). However, these cells did not express CD13 or high levels of KDR. Together, these patterns indicate that this protocol promotes the development of an epicardial-derived vascular population, distinct from the fibroblast populations characterized in our previous study. Given this, we will refer to it as a vascular progenitor (VP) population throughout this manuscript.

**Figure 1.**
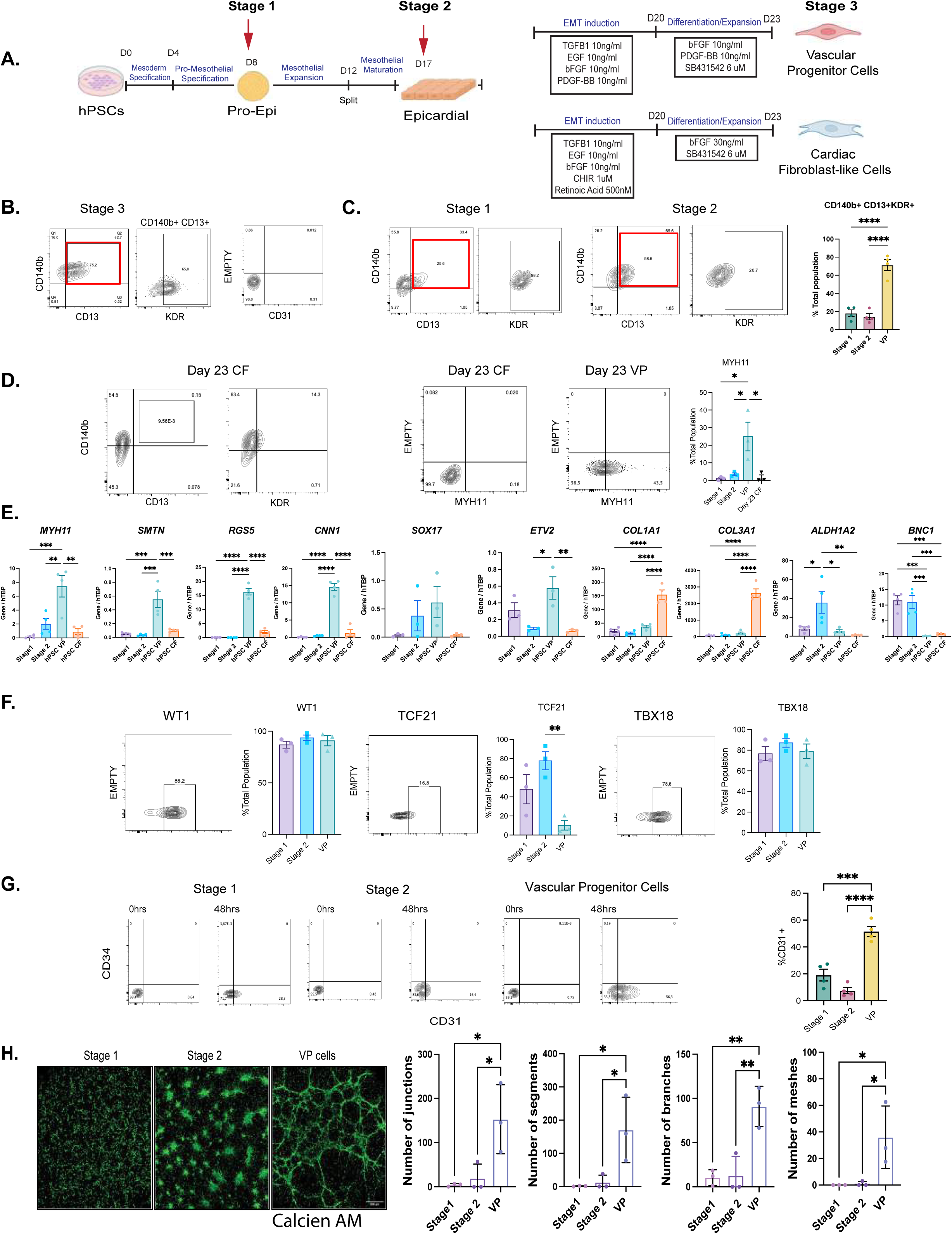
Generation of Epicardial-derived Vascular Progenitor Cells from Human Pluripotent Stem Cells. (A) Schematic of the staged differentiation protocol for generating vascular progenitor (VP) cells or cardiac fibroblast-like cells (CF) from human pluripotent stem cells (hPSCs) via an epicardial intermediate. The protocol proceeds through mesoderm specification (D0–D4), pro-mesothelial specification (D4–D8), mesothelial expansion (D8–D12), and mesothelial maturation (D12–D17; Stage 1 and Stage 2). From D17, EMT is induced using TGFB1 (10 ng/mL), EGF (10 ng/mL), bFGF (10 ng/mL), and PDGF-BB (10 ng/mL) through D20, followed by differentiation/expansion to D23 using either: bFGF (10 ng/mL), PDGF-BB (10 ng/mL), and SB431542 (6 μM) to generate Stage 3 Vascular Progenitor Cells, or bFGF (30 ng/mL), SB431542 (6 μM), CHIR (1 μM), and Retinoic Acid (500 nM) to generate Cardiac Fibroblast-like Cells. Cell icons created with BioRender.com (B) Representative flow cytometry plots of day 23 VP cells showing co-expression of CD140b and CD13 (left), KDR expression (middle), and absence of CD31 expression (right). KDR expression is measured from the CD140b^+^CD13^+^ gate, indicated by the red box. (C) Flow cytometric analysis of CD140b, CD13, and KDR expression on Stage 1 and Stage 2 epicardial populations (left two panels). Quantification of the proportion of CD140b^+^CD13^+^KDR^+^ triple-positive cells in the stage 1, stage 2, and VP populations (right bar graph). Error bars represent SEM. p < 0.05, **p < 0.01, ***p < 0.001, ****p < 0.0001 by one-way ANOVA with Tukey’s post-hoc test. (D) Flow cytometry analysis of CD140b, CD13 and KDR expression on day 23 cardiac fibroblast-like populations (CF; left panels), and intracellular flow cytometric analyses of MYH11 expression in day 23 CF and VP populations (middle, with empty control). Quantification of the per cent of MYH11^+^cells in the indicated populations (Bar graph; right) Error bars represent SEM. *p < 0.05 by one-way ANOVA with Tukey’s post-hoc test. (E) Quantitative RT-PCR analysis of expression of the smooth muscle/pericyte (*MYH11*, *SMTN*, *RGS5*, *CNN1*), endothelial/vascular transcription factors (*SOX17*, *ETV2*), extracellular matrix (*COL1A1*, *COL3A1*), and epicardial/mesothelial (*ALDH1A2*, *BNC1*), genes in the indicated populations. Expression is normalized to *hTBP*. Error bars represent SEM. *p < 0.05, **p < 0.01, ***p < 0.001, ****p < 0.0001 by one-way ANOVA with Tukey’s post-hoc test. (F) Intracellular flow cytometry analysis of expression of the epicardial transcription factors WT1, TCF21, and TBX18 in the indicated populations. Flow plots are shown only for the VP population. Graphs show quantification of the percentage of positive cells in each population. Error bars represent SEM. **p < 0.01 by one-way ANOVA with Tukey’s post-hoc test. (G) Left: Flow cytometric analysis of the CD31 and CD34 expression on the indicated populations prior to and following culture in the presence of VEGF (100 ng/ml) for 48 hours. Right: quantification of the % CD31+ cells in each population following 48 hours culture in VEGF. Error bars represent SEM. ***p < 0.001, ****p < 0.0001 by one-way ANOVA with Tukey’s post-hoc test. (H). Left: Representative fluorescence microscopy images of calcein AM-stained stage 1, stage 2, and VP cells following 48 hours of culture on Matrigel (left). Right: Quantification of network complexity parameters — number of junctions, number of segments, number of branches, and number of meshes — for each population (right bar graphs). Error bars represent SEM (n = 4 per group). *p < 0.05, **p < 0.01 by one-way ANOVA with Tukey’s post-hoc test.

Extended surface marker analyses of the stage 2 and stage 3 cells revealed dramatic changes consistent with a transition from an epithelial-like population (high E-CAD, EPCAM and ALDH), to a mesenchymal/vascular population defined by the upregulation of expression of CD73, CD105, CD44, CD200, NOTCH3, CD143, CD274, CD106, CD166 and reduction of epithelial markers (Figure S1A). The mesenchymal markers CD90 and PDPN were expressed on cells from both stages, a finding consistent with their known expression on both the epicardial and derivative populations (Mahtab E. et al., 2008). The dramatic changes in expression patterns of these markers between the two stages highlights the efficiency of the EMT with this protocol. Expression of CD73, CD105 and CD44 found on the MesoT population described by Colunga et al. (Colunga et al., 2019) and of CD274 used to identify human perivascular cells by Slukvin and colleagues (Kumar et al., 2017) suggests that our population shares characteristics with these previously described vascular cell types.

To further characterize the vascular nature of these cells, we carried out both molecular analyses and functional assessments of their smooth muscle and endothelial potential. RT-qPCR based analyses showed that the transition from the stage 2 population to the stage 3 VP cells was associated with an upregulation of expression of genes indicative of the smooth muscle lineage including *MYH11 SMTN*, *RGS5*, and *CNN1* (Figure 1E) and the early endothelial lineage including *SOX17* and *ETV2.* The levels of expression were significantly higher than those found in an epicardial-derived fibroblast population generated using our previously published protocol (Fernandes et. al., 2023). As expected, the fibroblasts expressed higher levels of *COL1A1* and *COL3A1,* functional genes associated with this lineage. The stage 1 and 2 cells had higher levels of *BNC1* and the stage 2 cells the highest levels of *ALDH1A2* consistent with expression patterns marking the epicardial lineage. In line with these expression patterns, intracellular flow cytometric analyses showed that the proportion of MYH11^+^ cells was significantly higher in the stage 3 than in the stage 2 population (Figure 1D).

To verify that these staged populations do indeed represent epicardial lineage development, we used intracellular flow cytometry to evaluate the expression patterns of transcription factors associated with specification, maturation and differentiation of this lineage *in vivo*. As shown in Figure 1F, the majority of cells at stage 2 (epicardial cells) showed elevated levels of WT1, TCF21, TBX18, proteins known to be expressed in epicardial cells *in vivo* (Martínez-Estrada et al., 2010; Cai et al., 2008; Acharya et al., 2012). This population also contained a high frequency of ISL1^+^, HAND2^+^ and NFATc^+^ cells. ISL1 marks the posterior second heart field progenitors from which the pro-epicardial lineage derives (Moretti et al., 2006; Cai et al., 2003), while HAND2 is expressed throughout the pro-epicardial organ and epicardium where it is required for EMT and coronary vessel formation (Barnes et al., 2011; Tsuchihashi et al., 2011). NFATc1 has been identified as a regulator of epicardial cell differentiation to derivative cell types (Combs & Yutzey, 2011). Together, the co-expression of these transcription factors confirms that this protocol recapitulates epicardial development (Figure S1B). Expression of WT1 and TBX18 along with the other transcription factors was maintained in the VPs, clearly demonstrating that these progenitors represent the epicardial lineage. In contrast to the patterns of these markers, the proportion of cells expressing TCF21, a transcription factor involved in the specification of epicardial cells to the fibroblast lineage (Acharya et al., 2012) declined significantly by stage 3, an observation consistent with specification to the vascular rather than the fibroblast lineage. GATA4, which plays a role in mesothelial specification (Watt et al., 2004), showed significantly higher expression in Stage 2 compared to Stage 1, consistent with its function during epicardial maturation. GATA3 expression was low across all stages, confirming the absence of contamination from intermediate mesoderm (renal/urogenital) lineages (Takasata et al., 2015). These findings confirm that this protocol efficiently promotes development most consistent with the human epicardial lineage as documented by the higher frequency of cells expressing the key transcription factors that regulate this lineage.

The above findings indicate that our modified protocol promotes the efficient specification of a population with vascular potential. As a functional assessment of smooth muscle potential of the stage 3 cells, we evaluated their response to endothelin-1 (ET-1). The response was assessed with rapid calcium flux and compared to that of staged matched epicardial-derived fibroblasts. As shown in Figure S1C treatment with 500 nM ET-1 elicited a robust and transient increase in intracellular calcium in the stage 3 cells but not in the fibroblasts, indicating that cells within the population display characteristics of vascular smooth muscle-competent progenitors

The observation that a subset of the day 23 VP population expresses KDR but not CD31 or CD34 suggests that it contains progenitors of the endothelial lineage. To further investigate this potential, we treated the cells for 48 hours with VEGF and then analyzed the population for expression of CD31 and CD34. Stage 1 and 2 cells were included for comparison. As shown in Figure 1G, the stage 3 VP cells did respond to VEGF and gave rise to a population that contained up to 50% CD31^+^ cells within 48 hours. Stage 1 and 2 did not respond as well and generated populations with significantly lower frequencies of CD31^+^ cells. Expression of CD34 was not upregulated.

As a measure of vascular potential, we next evaluated the capacity of the three different populations to form vascular networks *in vitro* following 24-hours of culture on Matrigel. The resulting structures were visualized using Calcein AM live-cell imaging. The outcome of this assay showed that the stage 3 VP cells, but not the earlier populations formed extensive, highly organized vascular networks with complex branching architecture. Quantitative analysis confirmed that the stage 3 cells generated significantly more junctions, segments, branches, and complete meshes compared to the other 2 populations. Taken together, these findings strongly support our interpretation that the stage 3 cells do, indeed represent a VP population.

### Engraftment potential of the VPs

As a first approach to assess the engraftment potential of the day 23 epicardial-derived VP population, we transplanted single cell suspensions into the mammary fat pad of NSG mice (Figure S2A). This site was chosen as multiple cell populations/mixtures can be quickly engrafted in each recipient with good host vascularization. Furthermore, grafts can be easily visualized and excised for analyses. In the first experiments, 1x10^6^ cells were injected into multiple fat pads. Four weeks later, the grafts were harvested, processed and tissue sections analyzed by immunohistochemistry for the presence of Ku80^+^ human cells and CD31^+^ human endothelial cells. To enable fluorescent tracking of transplanted cells, VP cells were derived from hPSC lines constitutively expressing tandem-dimer2, a red fluorescent protein (RFP) prior to transplantation (Irion et al., 2007). As shown in figure 2A, the cells did indeed engraft this site and formed well-organized vascular networks with numerous capillary-like structures. Notably, many of these structures had lumens containing mouse erythrocytes indicating that they had anastomosed with the host vasculature and were perfused. These findings strongly suggest that these structures represent functional vessels.

**Figure 2.**
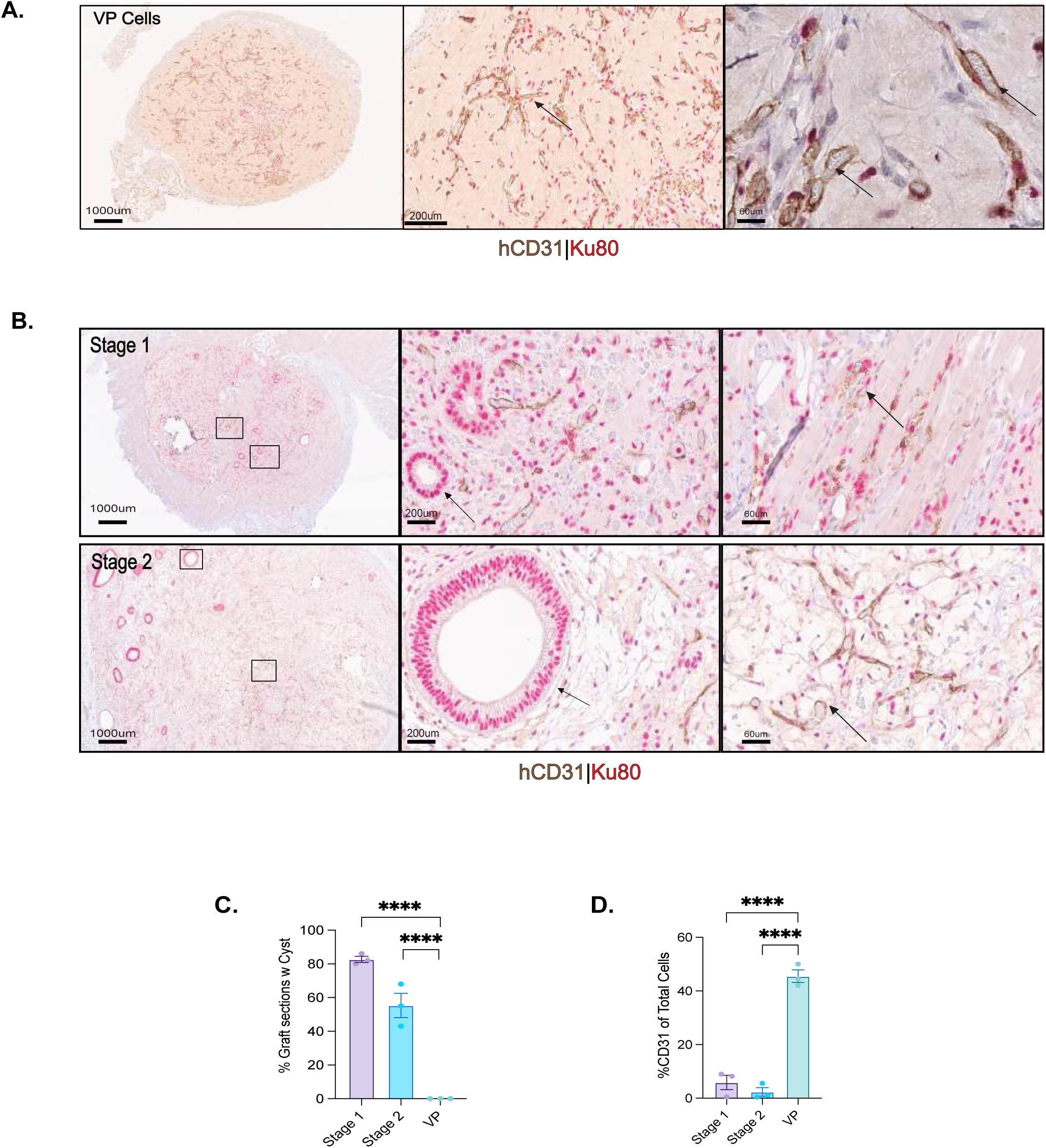
Engraftment potential of the VPs. (A) Representative immunohistochemistry images showing human CD31^+^ and Ku80^+^ cells in VP-derived grafts in fat pad tissue 28 days following transplantation of 1x10^6^ progenitors. Shown are low (left; scale bar = 1000 μm), medium (middle; scale bar = 200 μm), and high (right; scale bar = 40 μm) magnification views. Arrows indicate human CD31^+^ vascular structures embedded within the graft. (B) Representative immunohistochemistry images showing human CD31^+^ and Ku80^+^ cells in grafts generated from stage 1 or stage 2 cells 28 days following transplantation.1x10^6^ cells were transplanted. Low (left; scale bar = 1000 μm), medium (middle; scale bar = 200 μm), and high (right; scale bar = 60 μm) magnification views are shown. Arrows indicate representative structures. (C) Quantification of the percentage of the day 28 grafts generated from the indicated populations that contain cystic structures. Error bars represent SEM. ****p < 0.0001 by one-way ANOVA with Tukey’s post-hoc test. (D) Quantification of the percent of human Ku80^+^ cells that express CD31 in the day 28 grafts generated from the indicated populations. Error bars represent SEM. ****p < 0.0001 by one-way ANOVA with Tukey’s post-hoc test.

Given the robust engraftment potential of the day 23 population, we next wanted to determine if the earlier staged populations could generate comparable or perhaps better vascular grafts. The goal here was to identify the population with optimal engraftment potential. As shown in Figure 2B, Stage 1 and Stage 2 populations gave rise to identifiable grafts; however, these differed dramatically in composition from VP-derived grafts. Stage 1 and Stage 2 grafts were characterized by distinct cystic epithelial structures interspersed throughout regions of Ku80^+^CD31^−^ cells, which comprised the majority of the graft area. Quantification confirmed that the majority of grafts generated from Stage 1 and Stage 2 populations contained these cystic structures, whereas cystic tissue was absent from all VP-derived grafts (Figure 2C). Assessment of endothelial engraftment, measured as the proportion of Ku80+ human cells that co-expressed CD31, revealed that Stage 1 and Stage 2 grafts contained a low frequency of CD31+ cells (less than 10% of total Ku80^+^ human cells). In contrast, VP-derived grafts showed a striking increase in endothelial differentiation, with up to 50% of Ku80^+^ human cells expressing CD31 (Figure 2D). As indicated above, these CD31^+^ cells formed well-organized, vessel-like structures that were distributed throughout the graft.

As an independent measure of donor cells, we evaluated the expression of RFP in the different grafts, using a RFP specific antibody. The outcome of these analyses confirmed the presence of hPSC-derived grafts from each of the three populations and confirmed the differences in composition described above (Figure S2B).

### Optimization of VP fat pad engraftment

As a first step to optimize vascular engraftment in the fat pad, we tested the effect of VP cell dose on graft composition. The resulting grafts were harvested at day 28, processed and then analyzed quantitatively for the frequency of CD31^+^ cells per graft and the number of lumenized vessels per mm^2^. Increasing the number of cells transplanted from 1x10^6^ to 5x10^6^ resulted in a significant increase in the frequency of CD31^+^ cells detected and a 3-fold increase in vessel density in the grafts (Figure 3A). A further increase to 10x10^6^ cells per transplant had little effect and did not further improve the vascular composition of the grafts. Based on these findings, we used 5x10^6^ cells as the transplant dose for many of the following experiments.

**Figure 3.**
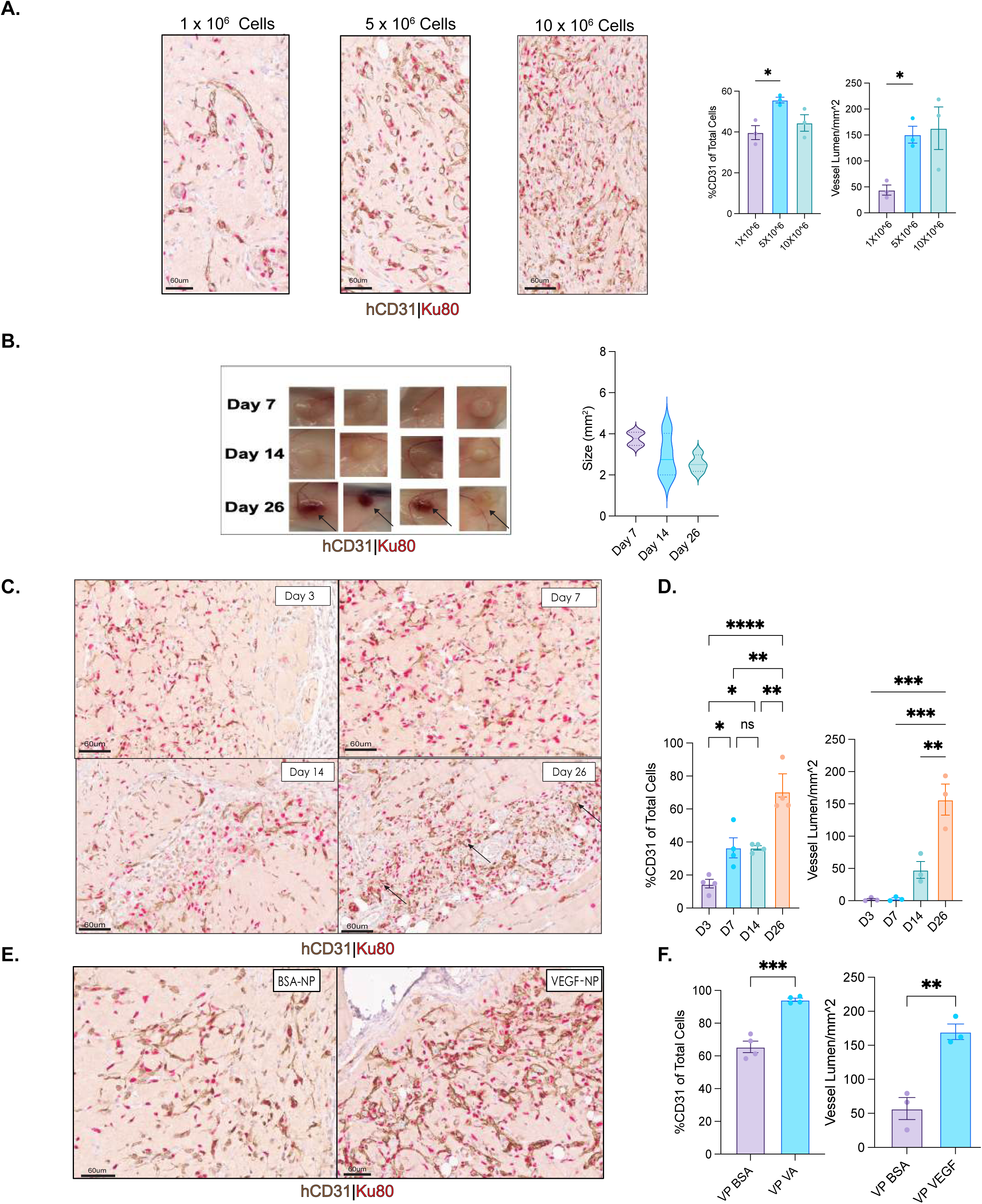
Optimization of VP fat pad engraftment. (A) Left: immunohistochemical analyses of CD31 and Ku80 expression in day 28 fat pad grafts generated with the indicated number of VPs. (scale bars = 60 μm). Right: Quantification of the percent of human Ku80^+^ cells that express CD31 and vessel lumen density in respective grafts. Error bars represent SEM. *p < 0.05 by one-way ANOVA with Tukey’s post-hoc test. (B) Left: Photos of intact fat pad grafts on the indicated day following transplantation of 5X10^6^ VPs. Arrows at day 26 indicate areas of visible vascularization. Right: violin plot analyses of quantification of graft size (mm²) at days 7, 14, and 26. (C) Representative immunohistochemical analyses of CD31 and Ku80 expression in fat pad grafts at days 3, 7, 14, and 26 following transplantation of 5x10^6^ VPs (scale bars = 60 μm). Arrows at day 26 indicate human CD31^+^ vessel-like structures. (D) Left: Quantification of the percent of human Ku80^+^ cells that express CD31 in day 3, 7, 14 and 26 grafts. Right: Quantification of the percent of human Ku80^+^ cells that express CD31 and vessel lumen density (vessel lumen/mm²) in the grafts at the indicated time points. Error bars represent SEM. *p < 0.05, **p < 0.01, ***p < 0.001, ****p < 0.0001 by one-way ANOVA with Tukey’s post-hoc test; ns = not significant. (E) Representative immunohistochemical analyses of CD31 and Ku80 expression in day 28 fat pad grafts generated from 1x10^6^ VPs transplanted together with either BSA loaded nanoparticles (BSA-NP) or VEGF-loaded control nanoparticles (VEGF-NP). scale bar = 60 μm. (F) Quantification of the percent of human Ku80^+^ cells that express CD31 (left) and the vessel lumen density (vessel lumen/mm²) (right) in grafts generated with VP cells together with the indicated nanoparticles. Error bars represent SEM. **p < 0.01, ***p < 0.001 by unpaired t-test.

All of the above analyses were carried out on grafts isolated 28 days following transplantation. We were next interested in characterizing the temporal dynamics of graft formation, specifically focussing on earlier timepoints (days 3, 7, and 12 in addition to 26) to establish the patterns of CD31^+^ cell maturation and vessel development. Gross examination of transplant sites showed the presence of comparable sized grafts at days 7, 14, and 26 post-transplantation (Figure 3B). Day 3 grafts were not included in this stage of analysis. The major difference between the grafts was the obvious progressive vascularization of those at day 26 (as shown by black arrows). Immunohistochemical analysis revealed that grafts could be detected within 3 days of transplantation (Figure 3C). At this stage, the grafts consisted of disperse groups of cells with minimal organization and relatively low frequencies (<20%) of CD31^+^ cells (Figure 3D). The frequency of CD31^+^ cells increased by days 7/14 and then again by day 26. Vessel structures were not detected in the early grafts but were present by day 14. Their frequency increased by more than 3-fold over the next 2 weeks. These patterns show that the VPs engraft rapidly and mature to CD31^+^ endothelial cells that form perfused vessels within 2 weeks of transplantation. The grafts continue to mature over the next 2 weeks as demonstrated by the increase in vessel content.

The extent of vascular development in the VP-derived grafts is likely influenced, in part, by the environment of the fat pad niche. To determine if it is possible to manipulate this environment and enhance vascular engraftment, we transplanted poly(lactic-co-glycolic acid) (PLGA) nanoparticles loaded with VEGF-A (VEGF-NP) together with the VPs into the fat pad. These particles release the factor in a sustained manner over a period of weeks, as characterized by *in vitro* release assays (data not shown). For these studies, a limiting lower dose (1x10^6^ cells) of VP cells was transplanted to increase the ratio of particles per target cell. We chose VEGF for the first analyses as the VEGF/KDR signaling pathway is known to promote vascular development and endothelial survival (Carmeliet et al., 1996; Gerber et al., 1998) and the VPs express KDR (Figure 1B). Nanoparticles loaded with BSA (BSA-NP) were transplanted as controls. As shown in Figures 3E and 3F, co-delivery of the VEGF-NPs significantly enhanced endothelial cell development in the grafts as demonstrated by higher frequencies of CD31^+^ cells and higher numbers of vessel structures compared to grafts initiated with the BSA-NPs. These findings show it is possible to promote vascular development within the grafts by manipulation of the environment through nanoparticle-based delivery of signaling pathway agonists.

Immunofluorescence analysis of day 26 grafts confirmed the maturation and functional organization of VP-derived vascular structures (Figure S3A-D). Co-staining for α-smooth muscle actin (ASMA) and CD31 revealed association of ASMA-positive mural cells with CD31-positive endothelial structures (Figure S3A). Analysis of CD144 (VE-cadherin) and MYH11 demonstrated endothelial junction formation within vessels surrounded by MYH11-positive smooth muscle cells (Figure S3B). Co-staining for CD140b (PDGFRβ) and CD31 confirmed the presence of CD140b-positive pericytes in close apposition to CD31-positive endothelial cells (Figure S3C). The CD140b-positive cells not in association with vessel structure may be fibroblast/mesenchymal derivatives. To assess arterial/venous vascular specification, we performed immunofluorescence for the human nuclear marker Ku80 together with EphB4 (vein) and EphB2 (artery) (Figure S3D). Human-derived cells expressing both ephrin receptors were identified within the grafts, indicating arteriovenous specification of VP-derived vessels. The maturation of these donor-derived networks was further characterized using human-specific CD144 (VE-cadherin) to visualize endothelial junctions (Figure S3E). Co-staining with mouse-specific CD31 demonstrated that the human vessels developed in close proximity to the endogenous host vasculature, confirming the formation of a chimeric vascular bed within the fat pad niche (Figure S3E).

To confirm the reproducibility of the VP cell differentiation protocol, we applied it to multiple independent hPSC lines (Figure S3E). Flow cytometric analysis of day 23 cells from HES2-tdRFP, ESI17, CHOP17, and PGPC17-11, lines demonstrated efficient generation of CD140b^+^CD13^+^KDR^+^ populations (55% to 80%) from all hPSC lines tested. Transplantation studies showed that the day 23 populations from 2 of the lines (ESI-17 and CHOP-17) generated vascular grafts in the fad pad site. These findings clearly demonstrate that this differentiation protocol is broadly applicable for the generation of transplantable VPs form different hPSC lines.

### VP engraftment of uninjured skeletal muscle

Having documented VP vascular engraftment in the fat pad, we were next interested in assessing the engraftment potential of these progenitors in a therapeutically relevant tissue, the hind limb skeletal muscle. For the first of these experiments, 5x10^6^ VPs were transplanted into the gastrocnemius muscle (hindlimb) of uninjured NSG mice (Figure S4A). Analyses of the transplanted tissue 4 weeks later showed the presence of extensive CD31^+^ vascular grafts that often covered the entire muscle tissue from patellar head of the vastus intermedialis of the quadricep to the femoral head (Figure 4A). Perfused vessel-like structures, similar to those found in the fat pad grafts were readily detected indicating that the cells could engraft and generate functional vasculature in skeletal muscle tissue.

**Figure 4.**
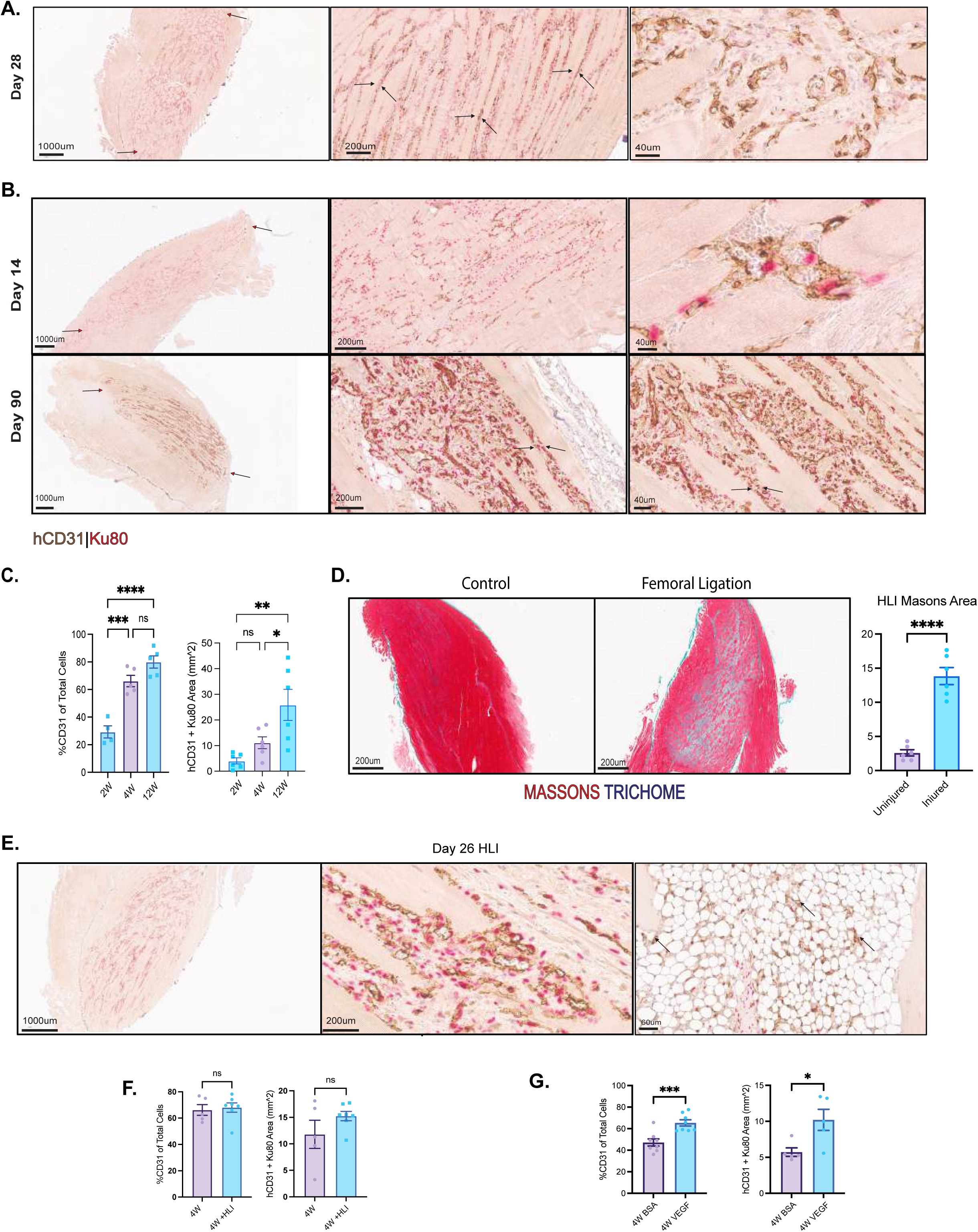
VP engraftment of uninjured skeletal muscle. (A) Representative immunohistochemical analyses of CD31 and Ku80 expression in day 28 skeletal muscle grafts generated from transplanted VPs (5X10^6^). Shown are low (scale bar = 1000 μm), medium (200 μm), and high (40 μm) magnification images. Arrows indicate representative graft boundaries in low magnification image. (B) Representative immunohistochemical analyses of CD31 and Ku80 expression in day 14 (top row) and day 90 (bottom row) skeletal muscle grafts generated from transplanted VPs (5X10^6^). Shown are low (scale bar = 1000 μm), medium (200 μm), and high (20–40 μm) magnification images. Arrows indicate human CD31^+^ structures. (C) Quantification of the percent of human Ku80^+^ cells that express CD31 (left) and of the total CD31^+^Ku80+ vascular area (mm²; right) in grafts at days 14, 28 and post-transplantation. Error bars represent SEM. *p < 0.05, **p < 0.01, ***p < 0.001, ****p < 0.0001; ns = not significant; one-way ANOVA with Tukey’s post-hoc test. (D) Representative immunohistochemistry showing masson’s trichrome staining of uninjured (control) and femoral ligation-injured hindlimb muscle (scale bars = 200 μm). Blue staining indicates injured fibrotic tissue. Quantification of fibrotic area (HLI Masson’s Area) in the control and injured tissue. Error bars represent SEM. ****p < 0.0001 by unpaired t-test. (E) Representative immunohistochemical analyses of CD31 and Ku80 expression in day 26 hindlimb ischemia (HLI) muscle grafts generated from transplanted VP cells (5x10^6^). Shown are images at low (1000 μm), medium (200 μm), and high (60 μm) magnification. Arrows indicate human CD31^+^ vascular structures engrafted within the ischemic adipose tissue. (F) Quantification of the percent of human Ku80^+^ cells that express CD31 (left) and of the total CD31^+^Ku80^+^ vascular area (mm²; right) in day 26 control and injured (HLI) muscle tissue. Error bars represent SEM; ns = not significant; unpaired t-test. (H) Quantification of the percent of human Ku80^+^ cells that express CD31 (left) and of the total CD31^+^Ku80^+^ vascular area (mm²; right) in day 28 ischemic skeletal muscle grafts generated with 1X10^6^ VPs transplanted with either BSA-loaded control nanoparticles (BSA-NP) or VEGF-loaded nanoparticles (VEGF-NP) at 4 weeks in hindlimb muscle. Error bars represent SEM. *p < 0.05, ***p < 0.001 by unpaired t-test.

To document the temporal patterns of vascular development in the hind limb muscle, we extended the analyses to include an earlier (14 days) and a later (90 days) time point post-transplant. As shown in Figure 4B, grafts were found dispersed throughout the transplanted tissue at all times analyzed. The grafts at day 90 remained large as shown by the presence of hCD31^+^/Ku80^+^ cells across the entire transplanted muscle tissue and contained numerous vessel-like structures. To assess changes in the grafts over the 90-day time period, we quantified the proportion of Ku80^+^ cells that express CD31 as in the above analyses of the fat pad grafts. Additionally, we also measured the total number of hCD31^+^Ku80^+^ cells in each graft as an indication of graft size (mm^2^) (Figure 4C). At the 2-week timepoint, a low frequency of human CD31-positive cells was detected dispersed throughout the graft. The frequency increased significantly over the next 2 weeks and then remained relatively constant for the 90-day duration of the study. Quantification of graft size revealed an increase over time, with significant differences observed between days 14 and 90 and 28 and 90.

At both days 28 and 90, the human vasculature appeared to align with the myofibrils of the mouse muscle tissue, with this alignment most prevalent in 90-day-old grafts, at which stage it can be easily visualized in low-power magnification images. Notably, this pattern of longitudinal vessel-to-myofiber alignment recapitulates the architecture of the native murine skeletal muscle vasculature, in which endomysial capillaries run parallel to myofibers and form organized microvascular units supplying them at regular intervals (Emerson & Segal, 1997; Hudlicka, 2011).

Vascular chimerism was also evident in the hindlimb musculature. Immunofluorescence for human CD144 and mouse CD31 revealed that donor-derived vessels integrated into the interstitial spaces between myofibers, often found in immediate association with expanded host microvessels (Figure S4B)

### VP engraftment in ischemic muscle

The above studies clearly show that the VPs can form vascular grafts in uninjured fat pad and skeletal muscle tissues. To determine if these progenitors can also engraft damaged tissue, we transplanted the cells into ischemic muscle tissue injured using two different approaches. In the first, mild ischemia was induced in the vastus intermedialis muscle through permanent femoral artery ligation (Figure S4C). Seven days following injury, 5x10^6^ VPs were transplanted into the damaged muscle. Uninjured muscle was transplanted as a control. Masson’s trichrome staining showed substantial areas (up to 13% of the tissue) of fibrosis in the damaged muscle compared to the control (<3% of tissue), demonstrating that femoral artery ligation did indeed induce the desired injury (Figure 4D). Analyses at 4 weeks post-transplantation revealed the presence of vascularized grafts in both the injured and control muscle. In the injured muscle, the grafts formed vascular networks in both the fibrotic and adipose regions of the damaged tissue (Figure 4E). Quantification of the grafts in both the ischemic and normal muscle showed that they contained comparable frequencies of CD31^+^ cells and were of similar size clearly demonstrating that the VPs can efficiently engraft the fibrotic microenvironment generated by ischemic injury (Figure 4F).

To assess whether VP engraftment could be augmented in damaged muscle, VP cells (1×10□) were co-transplanted with VEGF-loaded or BSA-loaded control nanoparticles into ischemic hindlimb and analyzed at 4 weeks. VEGF-NP co-delivery significantly increased the frequency of CD31^+^ cells within grafts relative to BSA-NP controls (Figure 4G, S4D). Consistent with this, quantification of total human vascular area (hCD31^+^Ku80^+^) confirmed that VEGF-NP-treated grafts contained significantly greater vascular tissue content, demonstrating that localized VEGF delivery enhances VP-derived vascular network formation even in the ischemic setting (Figure 4G).

Having established that VPs can engraft fibrotic ischemic tissue, we next sought to evaluate the therapeutic efficacy of this newly generated vasculature. To address this, we used a model of severe hindlimb ischemia induced by complete femoral artery resection (Figure S5A). This experimental procedure results in extensive ischemic injury and in most cases a loss of limb function. One week following injury, VP cells (5×10□) were transplanted into the medial aspect of the gastrocnemius and soleus of the injured limb. Matrigel without cells was transplanted as a control. In our initial study, mice were analyzed over a 3-week period using doppler imaging to monitor perfusion recovery in the injured limb (Figure S5B). Representative images at day 0 (post-surgery, pre-treatment) document perfusion deficits as visualized by the loss of high intensity flux signal in the ischemic limb. Quantification showed that VP transplanted animals recovered significantly greater levels of perfusion compared to Matrigel controls at 3 weeks (p=0.046) posttransplant.

Immunohistochemistry of extensor digitorum longus and adductor longus muscles confirmed VP cell engraftment within the ischemic tissue as demonstrated by the presence of human CD31^+^Ku80^+^ vascular structures integrated within the muscle architecture (Figure S5C). Vessel-like structures containing erythrocytes were readily detected in these grafts. These preliminary findings provide proof-of-concept that VP cell therapy improves perfusion recovery in severe hindlimb ischemia.

Given these findings, we next carried out more comprehensive efficacy studies with extended analyses to evaluate both perfusion recovery and functional outcomes (Figure 5A). For these studies, the injured muscle was transplanted with either VP cells (VP), VP cells together with VEGF-containing nanoparticles (VP+VEGF) or Matrigel alone (control). Cohorts of transplanted animals were followed for either 5 weeks or 3 months using laser Doppler perfusion imaging to monitor blood flow.

**Figure 5.**
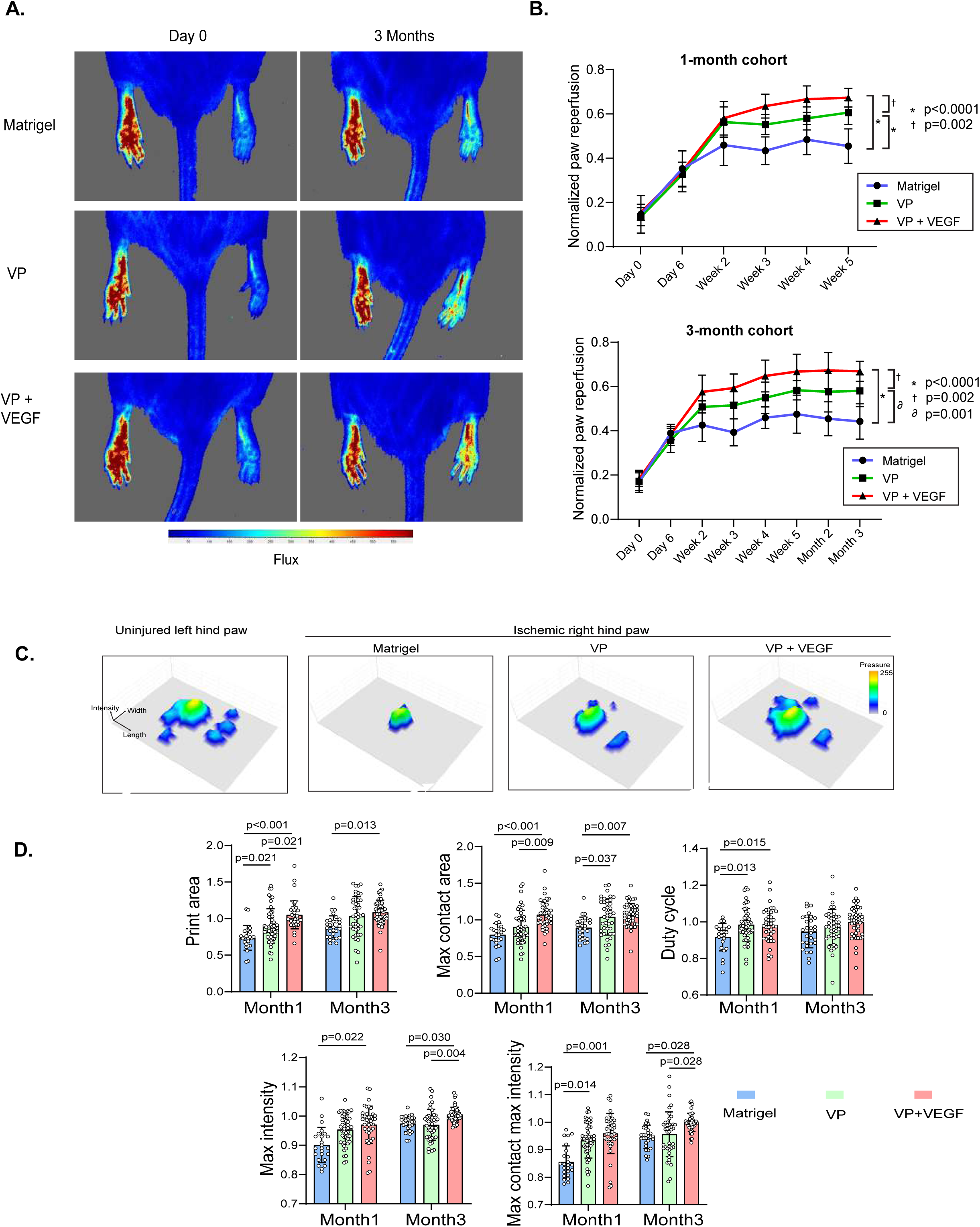
VP engraftment in ischemic muscle. (A) Representative laser Doppler perfusion images of NSG mice at day 0 (immediately post-ligation) and 3 months post-transplantation for Matrigel control, VP, and VP+VEGF nanoparticle treatment groups. Color scale indicates blood flow flux (blue = low, red = high). (B) Longitudinal quantification of normalized paw reperfusion (ischemic:contralateral limb ratio) for the 1-month cohort (top) and 3-month cohort (bottom). Error bars represent SEM * p < 0.0001 (VP or VP+VEGF vs. Matrigel); † p = 0.002 (VP+VEGF vs. Matrigel); ∂ p = 0.001 (VP+VEGF vs. VP); two-way ANOVA with Šídák’s multiple comparisons test. (C) Representative CatWalk pressure map images of the uninjured left hindpaw and ischemic right hindpaw from Matrigel, VP, and VP+VEGF-treated animals. Each 3D surface plot depicts print intensity (height), width, and length during the stance phase. Pressure scale: 0–255. (D). Quantification of weight-bearing endpoints (print area, maximum (Max) contact area, duty cycle, maximum intensity, maximum contact maximum intensity) at 1 and 3 months after intervention. n=24, 49, and 41 runs from 6, 9, and 9 mice at 1 month; n=29, 42, and 45 runs from 6, 9, and 9 mice at 3 months. Error bars represent SEM. Comparisons were made using a generalized linear mixed model. *P* < 0.05 are indicated.

In the first set of animals, both the VP and VP+VEGF transplanted animals showed significant improvement in perfusion at the 5-week time point compared to the controls (Figure 5B). Notably, perfusion in the VP+VEGF treated muscle was even better than that in the muscle that received VPs only, indicating that it is possible to manipulate the ischemic transplant environment with additional factors to increase efficacy of cell transplantation. The therapeutic benefit of VP transplantation was sustained to the 3-month timepoint (Figure 5A, B). As observed in the 5-week cohort, animals transplanted with either VP cells or VP cells plus VEGF showed significantly better levels of perfusion at 3 months compared to the control animals. Once again, transplantation of VEGF nanoparticles together with VPs led to significantly better recovery of perfusion than the VPs alone.

To determine whether improved perfusion translates to functional recovery, we next performed quantitative gait analysis using the CatWalk system, which measures weight-bearing capacity during ambulation (Figure S5D). For these analyses, the floor contacts of each paw were captured as diagonal gait images (Figure S5D). Three-dimensional paw print depictions prepared from the images were used to visualize and quantify the weight exerted on each paw (Figure 5C). These images showed that the ischemic right hind paw of the Matrigel-treated animals displayed reduced print intensity compared to the contralateral limb. The prints of the ischemic paws of the VP and VP+VEGF-treated animals, by contrast, showed greater intensity suggestive of some functional recovery. Videos of these animals demonstrated that the VP and VP+VEGF-treated groups exhibited improved limb placement and reduced limb drag during ambulation compared to Matrigel-treated controls, consistent with partial restoration of motor function in the ischemic limb (Supplemental Videos 1-3).

Quantitative analysis of weight-bearing endpoints at 1- and 3-months post-treatment demonstrated significant functional improvements following VP cell transplantation (Figure 5D). Print area was significantly increased in VP and VP+VEGF-treated animals at 1 month and in the VP+VEGF-treated animals at 3 months compared to controls. Maximum contact area showed significant improvements in VP+VEGF-treated animals at 1 month and in both the VP and VP+VEGF treated animals at 3 months. Duty cycle showed improvement in both transplanted groups at 1 month but not at 3 months. Maximum intensity measurements demonstrated improvements in the VP+VEGF-treated animals over the controls at 1 and 3 months and over the VP treated animals at 3 months. The VP only group also showed improvement over the control at the 1-month time point. Maximum contact maximum intensity similarly demonstrated improvements in the VP+VEGF-treated animals over the controls at 1 and 3 months and over the VP treated animals at 3 months. The VP only group also showed improvement over the control at the 1-month time point. Together, these comprehensive functional assessments demonstrate that VP cell therapy not only restores tissue perfusion but translates to meaningful improvement in weight-bearing capacity of the ischemic limb, with benefits sustained through at least 3 months post-treatment.

To determine if the functional improvements detailed above correlate with engraftment of human vascular cells, we performed detailed histological analysis of ischemic muscle tissue at 1- and 3-months post-treatment. Confocal microscopy of longitudinal sections through gastrocnemius muscle from VP+VEGF-treated animals at 3 months revealed the presence of human vasculature (Figure 6A). The transplanted VPs formed a substantial vascularized graft at the injection site, adjacent to the gastrocnemius muscle. This graft contained numerous CD31^+^Ku80^+^ human vessels that formed vascular channels filled with red blood cells visible by autofluorescence. The presence of these perfused vessels provides additional evidence of functional anastomosis of the human vasculature with the host circulation. Examination of the muscle tissue adjacent to the injection site revealed VP cell-derived microvessels (white arrows) within the interstitial space between skeletal myofibers (Figure 6A). Native mouse microvessels, visualized by mouse specific CD31 staining (yellow arrows), were also present throughout this region. In some instances, human and mouse vessels were found in close association, demonstrating vascular chimerism and integration of the VP-derived neovasculature with the host vascular network.

**Figure 6:**
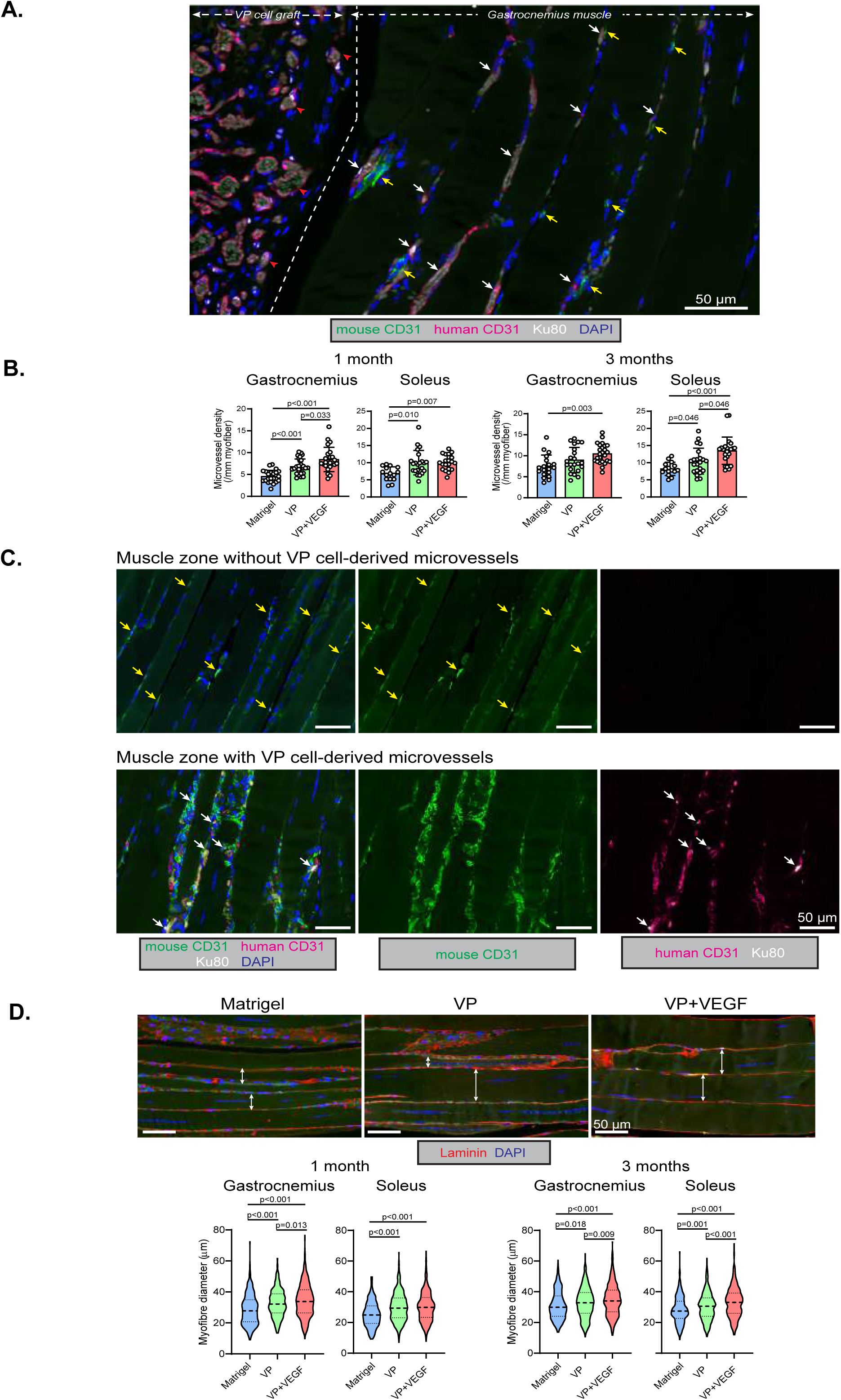
Histochemical analyses of transplanted ischemic hind limb muscle. (A) Confocal micrograph of a longitudinal section of ischemic mouse gastrocnemius muscle 3 months following injection of VP2 cells and VEGF-loaded nanoparticles. The tissue is immunostained for mouse CD31 (green), human CD31 (magenta), and Ku80 human nuclear antigen (white). Nuclei are stained with DAPI (blue). The injected VP cell - Matrigel mixture forms a heavily vascularized graft (left) adjacent to the gastrocnemius muscle (right), wherein all vascular endothelial cells are of human origin. Red blood cells can be seen by autofluorescence in the lumen of the vascular channels, suggesting functional anastomosis with the host circulation. Within the adjacent muscle, there is an abundance of VP-derived microvessels in the interstitial space between skeletal myofibers (white arrows depicting either the human CD31 signal and/or human nuclear antigen signal). Native mouse microvessels are also present (yellow arrows) and human and mouse vessels can be found closely associated. Scale bar = 50 μm. (B) Quantification of the density of mouse CD31+ microvessels in ischemia-injured gastronemius and soleus muscles, 1- and 3-months post-injection of the designated material. Differences were assessed using a generalized linear mixed model with the specific injection group as fixed factor and the individual mouse as a random factor. Multiple comparisons were accounted for using the sequential Sidak test. n=23, 28, and 23 regions of interest (ROIs) in 1-month gastrocnemius muscle, n=15, 22, and 20 ROIs in 1-month soleus muscle, n=19, 22, and 25 ROIs in 3-month gastrocnemius muscle, and n=15, 22, and 24 in 3-month soleus muscle, all from 6, 9, and 9 mice for the respective injection. (C) Confocal micrographs of ischemia-injured gastrocnemius muscle 3 months after injection of VP cells. Muscle regions without (upper panels) and with (lower panels) VP2 cell-derived microvessels are shown, with yellow and white arrows indicating mouse and human endothelial cells, respectively. Scale bar = 50 μm. (D) Confocal micrographs of ischemia-injured gastrocnemius muscle 3 months after injection of Matrigel only, VP cells, or VP cells and VEGF-loaded nanoparticles, immunostained for laminin (red) to delineate the myofiber boundaries. Nuclei were stained with DAPI (blue). Faint green autofluorescence was overlaid to visualize the skeletal myofibers and double-sided arrows depict representative myofiber diameters, which are variably increased in VP cell-injected muscles and homogeneously increased in VP2 cell and VEGF-loaded nanoparticle-injected muscles. Quantification of the diameters of individual myofibers 1- and 3-months post-injection. Differences were assessed using the Kruskal-Wallis test and Dunn’s post-hoc testing for multiple comparisons. n=300, 450, and 450 myofibers from 6, 9, and 9 mice, respectively. Scale bar = 50 μm.

Quantification of mouse CD31-positive microvessel density revealed that transplantation of either VPs alone or the combination of VPs and VEGF significantly increased the abundance of endogenous vessels in ischemic muscle (Figure 6B). At 1-month, both the gastrocnemius and soleus muscles showed increased endogenous microvessel density in both transplant groups compared to the control. At 3 months, these elevated levels were maintained in both muscle groups in the VP+VEGF treated animal, while the soleus muscle showed higher density in the animals that received only the VP cells. Beyond these differences, we also observed that the gastrocnemius muscle at 1 month and the soleus muscle at 3 months in mice that received VPs+VEGF had higher microvessel density than mice that were transplanted with VPs alone.

To determine if this angiogenic effect of the transplanted human cells on the mouse vasculature is localized to areas adjacent to the engrafted human vasculature or if it is observed more broadly, beyond the graft site, we compared muscle regions containing VP-derived microvessels to regions lacking human vascular cells within the same tissue sections (Figure 6C). In zones without VP cell-derived microvessels, mouse CD31-positive endogenous vessels were present at baseline levels. In striking contrast, zones containing VP-derived microvessels exhibited not only the expected human-derived vascular structures but also a markedly increased density of endogenous mouse CD31-positive microvessels. This observation indicates that VP cells both directly contribute to neovascularization through differentiation into functional endothelium and indirectly stimulate an endogenous angiogenic response in the surrounding tissue.

The increased vascularization was accompanied by preservation of muscle architecture, as assessed by myofiber diameter measurements (Figure 6D). Laminin immunostaining to delineate myofiber boundaries revealed substantial differences in myofiber size across treatment groups. Matrigel-treated ischemic muscle contained small, irregular myofibers characteristic of chronic ischemic injury. VP cell-treated muscle showed variably increased myofiber diameters, with some fibers approaching normal dimensions while others remained small. VP+VEGF-treated muscle demonstrated homogeneously increased myofiber diameters, with more uniform restoration of fiber size throughout the tissue. Quantitative analysis confirmed these observations (Figure 6D). At 1-month post-treatment, both gastrocnemius and soleus muscles showed significantly larger myofiber diameters in VP and VP+VEGF-treated animals compared to Matrigel controls, with VP+VEGF showing additional benefit over VP alone in the gastrocnemius. These differences were sustained at 3 months, with both VP and VP+VEGF groups maintaining significantly larger myofiber diameters than Matrigel controls in both muscle groups, and VP+VEGF showing improvement over VP alone in both gastrocnemius and soleus.

Together, these findings demonstrate that VP cell therapy provides therapeutic benefit through multiple mechanisms: direct contribution to the neovasculature through formation of perfused, host-integrated human vessels; paracrine stimulation of endogenous angiogenesis in surrounding tissue; and consequent preservation of myofiber integrity in chronically ischemic muscle.

### Single-Cell RNA-seq analysis of VPs and derivative endothelial cells

To gain further insight into the molecular characteristics of VPs and derivative endothelial populations, we performed single-nucleus RNA sequencing on the population used for transplantation and on cells isolated from fat pad and hindlimb grafts at 1-month post-transplantation (Figure 7). Nanoparticles were not co-injected for these studies. Integration of all three datasets revealed distinct transcriptional states corresponding to pre-transplant VP and post-transplant vascular populations from each tissue site.

**Figure 7.**
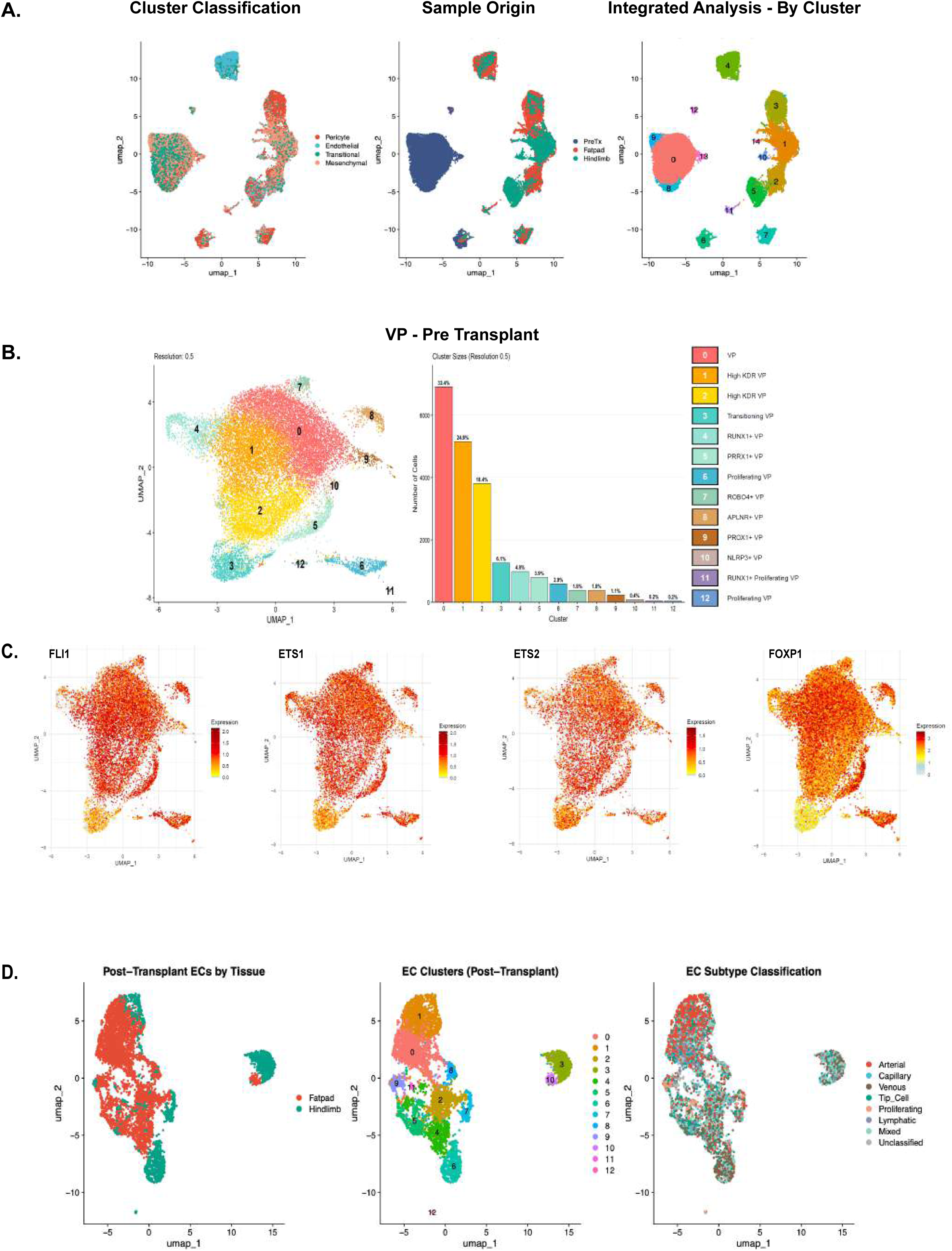
Single-Cell RNA-seq analysis of VPs and derivative endothelial cells. (A) UMAP visualization of the integrated single-cell RNA-seq dataset colored by cell type classification (pericyte, endothelial, transitional, mesenchymal; left), sample origin (pre-transplant VP, fatpad, hindlimb; middle), and cluster identity from integrated analysis (right), with numbered clusters labeled. (B) UMAP of pre-transplant VP cells at clustering resolution 0.5, with cells colored by cluster identity (left) and a bar graph (right) showing the proportional size of each cluster as a percentage of total pre-transplant VP cells. Cluster identities are indicated in the accompanying legend. (C) Feature plots showing expression of the vascular transcription factors *FLI1*, *ETS1*, *ETS2*, and *FOXP1* across all cells in the pre-transplant VP UMAP, with expression levels indicated by color scale (low = yellow, high = red). (D) UMAP visualization of post-transplant endothelial cells colored by tissue of origin (fatpad vs. hindlimb; left), EC cluster identity (middle), and EC subtype classification (right), including arterial, capillary, venous, tip cell, proliferating, lymphatic, mixed, and unclassified populations.

UMAP visualization showed transcriptionally distinct clusters, with pre-transplant VP cells forming pericyte, endothelial, transitional and mesenchymal clusters post annotation with module scores. Interestingly, post-transplant populations from fat pad and hindlimb occupied separate transcriptional territories (Figure 7A, middle panel) (Supplemental Table 1). An integrated analysis confirmed that the three sample origins were clearly resolved by cluster identity, with pre-transplant VP cells partially overlapping with both fat pad and hindlimb-derived populations (Figure 7A, right panel). This organization indicates that VP cells undergo tissue-specific transcriptional changes following each engraftment niche while retaining molecular relationships to their pre-transplant state.

Analysis of pre-transplant VP population alone identified 12 distinct clusters reflecting some molecular diversity within the population (Figure 7B). These clusters differed in their expression of genes associated with mesenchymal and vascular development. For instance, the largest clusters, clusters 0, 1 and 2 could be distinguished based on the expression of *CD140B* and *CD13* (cluster 0) or *KDR* (clusters 1 and 2). The remaining smaller clusters were characterized by elevated levels of *ETV2* (cluster 3), *RUNX1* (cluster 4), *PRRX1* (cluster 5), *ROBO4* (cluster 7), *APLNR* (cluster 8), *PROX1* (cluster 9), and *NLRP3* (cluster 10) and *MKI67* (clusters 6, 11 and 12). An example of the differential expression patterns of *PRRX1*, *ETV2* and *MKI67* as determined by feature plot analyses differences is shown in Figure S6A. While these differences document heterogeneity within the population, the broad expression of *FLI1*, *ETS1*, *ETS2*, and *FOXP1*, core transcription factors involved in the specification and function of the endothelial lineage together with the combined expression patterns of the above genes indicates that a large majority of the cells in the day 23 VP population express a transcriptional program consistent with vascular lineage competence (Figure 7C). The lack of CD34 and CD31 protein expression (Figure 1B) at this stage suggests that the endothelial lineage cells within the population represent progenitors not mature endothelium.

Violin plot analysis of key lineage markers across the 13 pre-transplant VP clusters confirmed that the day 23 VP population represents progeny of the epicardial lineage (Figure S6B). Broad expression of *WT1, TBX18, HAND2*, and *GATA4* across most clusters reinforces this conclusion, consistent with our earlier flow cytometric analyses. In contrast, *GATA3* and *ISL1* were restricted to clusters 9–11, identifying a transcriptionally distinct subpopulation, while *TCF21* and *NFATC1* showed sparse expression throughout. Broad expression of *PDGFRB* and *KDR* in most clusters further validates the signature surface phenotype of the population and corroborates our flow cytometric findings, although *ANPEP* (CD13), *ISL1* and *NFATc* transcript levels were low despite robust protein detection by flow cytometry (Supplemental Figure 1B).

For characterization of the VP-derived engrafted progeny we focused our analyses on populations within the integrated dataset that expressed endothelial markers. Feature plot analysis of the core endothelial markers *PECAM1* (CD31), *CDH5* (VE-cadherin), *KDR*, and *VWF* documented broad endothelial identity in both fat pad and hindlimb grafts, with *PECAM1* and *KDR* showing the widest expression and *CDH5* and *VWF* enriched in specific subpopulations. The patterns confirm the endothelial nature of the engrafted VP-derived progeny (Figure S6C).

Classification of post-transplant cells into endothelial, mesenchymal, and mixed populations revealed that both tissue sites contained all three cell types (Figure S6D). However, endothelial representation was low at both sites, with hindlimb grafts showing even poorer endothelial coverage than fat pad grafts. This likely reflects sampling bias inherent to single-cell preparations from these tissues, as endothelial cells are prone to underrepresentation following dissociation, particularly from skeletal muscle. The predominance of mesenchymal cells in both conditions may therefore reflect a recovery artifact rather than a true shift in VP cell fate.

Post-transplant endothelial cells from both tissue sites were visualized separately and as an integrated dataset (Figure 7D). UMAP analysis showed that endothelial cells from the fat pad and hindlimb occupied largely overlapping but distinguishable transcriptional territories. Sub-clustering of the combined endothelial populations identified 13 clusters (clusters 0–12). Dot plot analysis of gene expression across the endothelial clusters from each tissue site showed that core endothelial markers (*PECAM1*, *CDH5*, *KDR*, *CD34*) were broadly expressed across clusters from both fat pad and hindlimb (Figure S7A). However, the two sites showed distinct enrichment patterns. Fat pad endothelial clusters showed prominent expression of arterial markers *EFNB2*, *DLL4*, and *HEY1* particularly in clusters 9 and 1, whereas hindlimb clusters showed broader expression of venous markers *NR2F2* and *EPHB4*. Notably, *GJA5* and *SOX17*, markers of arterial identity, were enriched in specific clusters in both tissues. Proliferative markers (*MKI67*, *TOP2A*, *CCNB1*) were concentrated in discrete clusters at each site. Endothelial subtype annotation revealed transcriptional signatures indicative of the presence of arterial, capillary, venous, lymphatic, tip cell, proliferating and mixed populations (Figure 7D, right panel). Consistent with this, feature plot analysis confirmed the tissue specific distribution of arterial markers *EFNB2*, *DLL4*, and *HEY1*, venous markers *NR2F2* and *EPHB4*, and the capillary marker *RGCC* across the integrated endothelial UMAP, with arterial marker expression enriched in the fat pad-associated cluster and venous and capillary markers more broadly distributed across both tissue populations.

Analysis of signaling pathway activity revealed niche-specific transcriptional programs (Figure S7C). Module scoring for key signaling pathways demonstrated differential activity between fat pad and hindlimb endothelial populations. Fat pad-derived endothelial cells showed significantly higher activity for shear stress mechanotransduction, VEGF signaling, hypoxia-HIF, angiopoietin signaling, NOTCH signaling, and MAPK-ERK pathways. In contrast, hindlimb-derived endothelial cells were enriched for Hedgehog, non-canonical WNT, EGF, PDGF, FGF, TGFβ, canonical WNT, and retinoic acid signaling. The enrichment of VEGF and NOTCH signaling in the fat pad endothelium may reflect the highly vascularized and remodeling nature of adipose tissue (Sung et al., 2013; Corvera & Gealekman, 2014), while the enrichment of growth factor signaling pathways including FGF, PDGF, and EGF in the hindlimb is consistent with the regenerative microenvironment of skeletal muscle (Christov et al., 2007).

Examination of angiogenic ligand expression revealed tissue-specific patterns (Figure S7D). Dot plot analysis showed differential expression of VEGF family members (*VEGFA*, *VEGFB*, *VEGFC*, *VEGFD*), *PGF*, *IGF1*, *IGF2*, FGF family members (*FGF1*, *FGF2*, *FGF7*, *FGF10*), *JAG1*, *DLL4*, *ANGPT1*, and *ANGPT2* between fat pad and hindlimb samples. Among the most notable differences, fat pad-derived endothelial cells showed higher expression of *ANGPT1*, *ANGPT2*, and *PGF*, whereas hindlimb-derived cells were enriched for *IGF1 IGF2, VEGFB*, and *VEGFD* expression. The enrichment of Notch ligands *JAG1* and *DLL4* in the fat pad is consistent with the higher proportion of arterial subtypes observed in this tissue. Taken together, these findings indicate that the endothelial cells in the grafts from the two sites acquire tissue-specific molecular identities that reflect adaptation to the local niche.

## Discussion

The ability to revascularize injured and ischemic tissues through the generation of new blood vessels from transplanted progenitors represents a major advance towards novel therapies for peripheral vascular diseases. While the therapeutic potential of *de novo* vessel generation in injured tissue has been long recognized, the development of robust and efficacious cell therapy-based revascularization strategies has been challenging (Annex, 2013; Cooke & Losordo, 2015). This difficulty reflects, in part, the complicated multi-step process required to create a functional 3-dimensional blood vessel from a single cell suspension of progenitor cells. The key steps include engraftment and survival of the transplanted cells, differentiation of these cells to an endothelial cell fate, formation of vessels from the endothelial cells and finally integration of the new vessels with the host vasculature. In this study, we report on the generation of hPSC-derived progenitors that fulfill these criteria and display the ability to stably engraft both fat pad and skeletal muscle tissue of NSG mice where they form human vessels that functionally anastomose with the host vasculature. Notably, when transplanted into ischemic hind limb muscle, these progenitors generate new vasculature that significantly improves perfusion in the damaged tissue and partially restores limb function over a 3-month period. Together, these findings document the feasibility of revascularizing transplanted tissues in a pre-clinical model and in doing so represent an important step towards the development of novel cell-based therapies to treat ischemia.

A defining feature of the engraftable VP progenitors described here is their development from an epicardial intermediate, a cell type characterized by the co-expression of a core set of transcription factors including WT1, TCF1 and TBX18. The derivative VP population, which retains aspects of this epicardial signature and was initially identified by the co-expression of CD140b, CD13 and KDR, displays a range of cell surface markers and gene expression patterns (Figures 1-S1) associated with smooth muscle/pericyte and endothelial cell development. The broad expression patterns of a core set of endothelial transcription factors, *FLI1*, *ETS1*, *ETS2* and *FOXP1* indicates that the population is enriched in endothelial lineage cells, a finding that correlates well with its vascularization potential. Although differentiated over a period of 23 days, the VPs surprisingly have not progressed beyond the progenitor stage of development as demonstrated by the lack of CD34 and CD31 surface marker expression. These properties distinguish the VPs from most other hPSC-derived endothelial populations reported to date that are typically identified by the co-expression CD34, CD31 and/or CD144 and specified within several weeks of differentiation (Wang et al., 2020; Pan et al., 2024).

Our findings that the engraftable VPs develop from an epicardial-like intermediate are in line with those of Colunga et al. (Colunga et al., 2019) that described a hPSC mesothelial-derived endothelial/smooth muscle bi-potential progenitor (MesoT) that displays the capacity to vascularize a neonatal injured mouse heart. Although the protocol used for the generation of the MesoT progenitors differs somewhat from the one we designed for the VPs, the two derivative populations share some properties including the expression of the transcription factors WT1, TBX18, and TCF21 and the surface markers CD44, CD73 and CD105 as well as the ability to generate both smooth muscle and endothelial progeny. Comparison of the *in vivo* potential of the two populations, however, is not straightforward, as we focused our studies on extended engraftment of adult tissue whereas MesoT cells were evaluated in a neonatal heart injury model and in decellularized vascular scaffolds anastomosed to the host circulation, contexts that differ substantially from direct transplantation into adult ischemic tissue. The robust endothelial potential of the epicardial derived VPs is not predicted from studies on the mouse epicardium that collectively show that this population has limited endothelial potential *in vivo* (Cai et al., 2008; Acharya et al., 2012; Zhou and Pu, 2012). These apparent differences may reflect differences between mouse and human epicardial cells. Alternatively, the hPSC-derived epicardial-like population generated with our protocol *in vitro* may not be equivalent to the epicardium *in vivo*.

While the endothelium throughout our body shares expression of a core set of molecular markers that define the endothelial lineage, tissue specific differences are also well documented, reflecting the unique vascular demands of the different organs (Kaluka et al., 2020; Trimm and Red-Horse, 2023; Hou et al., 2022; Lin et al., 2026). The underlying developmental events that ultimately lead to the formation of these distinct vascular subtypes are not fully understood, however they most likely reflect contribution from distinct embryonic progenitors as well as patterning by the unique environments of the different tissues (Trimm and Red-Horse, 2023; Gomez-Salinero and Rafii, 2018; Lin et al., 2026). Given this, an important issue faced when designing a vascular therapy, is whether or not one progenitor population can engraft and vascularize different organs. While we did not carry out extensive multi-organ analyses, our findings clearly show that the VPs can vascularize both the fat pad and skeletal muscle. Notably, our molecular profiling showed that the resulting endothelial cells displayed some differences including in expression of components of the *TGF*β*, Wnt, PDGF, Notch, VEGF, FGF* and angiopoietin signaling pathways and in the proportion of capillary phenotypes generated in the 2 different sites. These findings suggest that VP cell derivatives can respond to local microenvironmental cues, acquiring tissue-appropriate characteristics.

Additional evidence that local environments can influence engraftment was provided by our studies showing that co-transplantation of VEGF containing nanoparticles significantly enhanced the extent of vascularization. These findings clearly demonstrate that it is possible to experimentally manipulate VP-mediated engraftment and pave the way for future studies aimed at identifying other factors that can impact engraftment and vascularization. Beyond local microenvironment influences that shape vascular development, different organs appear to display different levels of ‘permissiveness’ to vascular engraftment. This is best exemplified by transplants with the CD31^+^CD34^+^ endothelial population that engrafted but failed to generate numerous perfused vessels in the fat pad but, as documented in our previous studies, is able to efficiently engraft the liver and generate LSECs (Gage et al., 2020 and 2022). The underlying mechanisms that control site specific engraftment are currently unknown.

Successful cell-based therapy for peripheral vascular disease will likely require long-term revascularization of damaged, ischemic muscle tissue. Our studies showing extensive vascular engraftment in non-injured skeletal muscle over a 3-month period and the demonstration that engraftment in ischemic tissue partially restores perfusion and hind limb function, provides compelling evidence that VPs represent an attractive candidate population for this type of therapy. Together with restoration of blood flow, the hPSC-derived vasculature appeared to also preserve muscle architecture, demonstrating the desired beneficial effect of a cell-based therapy. The extensive levels of sustained vascular engraftment from transplanted single cell suspensions of VPs distinguishes the *in vivo* potential of these progenitors from that of most other hPSC-derived vascular cell types reported to date. For instance, Gong et al. recently demonstrated that transplantation of vascular organoids generated from endothelial and vascular smooth muscle cells specified through enforced transcription factor (ETV2, NKX3.1) expression into ischemic hind limb muscle could improve perfusion and reduced fibrosis in the tissue over a 2-week period (Gong et al., 2025). However, in contrast to the organoids, single cell suspensions of the endothelial and vascular smooth muscle cells did not engraft, demonstrating a striking difference in potential from our VPs. Given that the hind limb analysis in this study was only carried out for 2 weeks, it is unclear if the organoids could sustain engraftment and functional improvement over longer periods of time. Recently, Heuslein et al. described a 10-day differentiation protocol to produce vascular cells that were transplanted into ischemic hind limb muscle in immune compromised mice (Heuslein et al., 2026). Analyses of these animals revealed low levels of engraftment that declined within several weeks of transplantation. Although vascular engraftment was not well documented at any stage in these animals, the injured muscle did show improved perfusion over a 64-day period, suggesting a paracrine mechanism of action, possibly though the recruitment of host vasculature. We also observed an enhancement of host vasculature in the ischemic muscle, but in this case, it appeared to be restricted to regions of the graft that contained hPSC-derived cells, suggesting a localized effect. Taken together, the collective findings from these studies suggest that both donor-derived *de novo* vasculogenesis and host angiogenesis can improve specific functional properties of ischemic tissue. While both may be considered in the design of cell-based therapies for certain diseases, the development of new vasculature from transplanted progenitors would be the preferred approach for the treatment patients with compromised endothelial cell function.

In summary, we have developed a developmental biology-informed approach for generating vascular progenitor cells with unprecedented engraftment capacity. The staged differentiation through an epicardial intermediate, the requirement for this developmental trajectory for functional vessel formation, and the sustained therapeutic benefit in hindlimb ischemia with corresponding functional recovery, together provide a foundation for advancing cell-based revascularization therapy toward clinical application.

## Limitations of Study

Several limitations should be considered when interpreting these findings. First, all in vivo transplantation experiments were performed in immunodeficient NSG mice to enable engraftment of human cells. Additionally, mice have robust collateral vessel formation, unlike PAD patients. Clinical application would require evaluation of VP cell survival and function in the context of an intact allogeneic immune response, or alternatively, the use of autologous patient-specific iPSC-derived VP cells. Second, while therapeutic benefit was sustained through three months in the hindlimb ischemia model, longer follow-up studies spanning six to twelve months will be required to confirm the long-term stability of engrafted vessels in the context of chronic vascular disease. Third, our studies were conducted in otherwise healthy immunodeficient mice; whether VP cells retain their engraftment capacity and therapeutic efficacy in the presence of comorbidities such as diabetes, atherosclerosis, or advanced age, conditions prevalent in patients with peripheral arterial disease, remains to be investigated in detail. Fourth, single-cell transcriptomic profiling of engrafted cells was performed at a single post-transplant timepoint; longitudinal profiling would provide a more complete picture of the dynamics of endothelial specification and tissue-specific adaptation.

## Acknowledgments

We thank the Keller, Pickering, and Shoichet labs for their advice on the manuscript and experiments. We also thank the University Health Network/SickKids flow cytometry facility, UHN Pathology Research Program, the Princess Margaret Genomics Centre, and the Advanced Optical Microscopy Facility at the Princess Margaret Cancer Research Tower for their expert technical assistance. We acknowledge the support of the Government of Canada’s New Frontiers in Research Fund (NFRF), [NFRFT-2022-00447]. This work was partly supported by Bluerock Therapeutics. This work was supported through the Canadian Institutes of Health Research (G.M.K., FDN159937).

## Author Contributions

I.M.F. and G.M.K. conceived the project. I.M.F. designed and performed the differentiation protocol development, flow cytometry, fat pad and hindlimb transplantation, immunohistochemistry, scRNA-seq experiments and bioinformatic analysis, and wrote the manuscript. H.Y. and Z.N. performed the severe hindlimb ischemia model, laser Doppler perfusion imaging, CatWalk gait analysis, and associated immunohistochemistry under the supervision of J.G.P. Y.Y. designed and synthesized the VEGF-loaded PLGA nanoparticles and performed branching assays under the supervision of M.S. B.K.G. helped establish and performed fat pad transplantation experiments. M.G. assisted with cell culture and flow cytometry. G.M.K. supervised the project and acquired funding. All authors read and revised the manuscript.

## STAR Methods

### RESOURCE AVAILABILITY

#### Lead Contact

Further information and requests for resources and reagents should be directed to and will be fulfilled by the Lead Contact, Gordon Keller.

#### Materials Availability

All unique reagents generated in this study are available from the Lead Contact upon reasonable request.

#### Data and Code Availability

Single-cell RNA-sequencing data generated in this study have been deposited in GEO and are publicly available as of the date of publication. Accession numbers are listed in the Key Resources Table. Any additional information required to reanalyze the data reported in this paper is available from the Lead Contact upon request.

### EXPERIMENTAL MODEL AND SUBJECT DETAILS

#### Human Pluripotent Stem Cell Lines

hPSC lines (HES2-tdRFP/eGFP, ESI-17) were maintained on gelatin (0.1%w/v in PBS) and growth factor reduced Matrigel (0.25%v/v, Corning) coated dishes with irradiated mouse embryonic fibroblasts in hPSC culture media consisting of DMEM/F12 (Cellgro) supplemented with penicillin/streptomycin (1%, ThermoFisher), L-glutamine (2mM, Thermo- Fisher), non-essential amino acids (1x, ThermoFisher), b-mercaptoethanol (55mM, ThermoFisher), KnockOut serum replacement (20% v/v, ThermoFisher) and rhbFGF (10-20ng/ml optimized for each line, R&D).

iPSC lines (CHOP-17, PGPC17-11) were maintained feeder-free on tissue culture plates coated with 1.5% v/v growth factor-reduced Matrigel (Corning). Matrigel coating was prepared by diluting Matrigel to 1.5% (v/v) in cold IMDM, applying to plates, and incubating for 1 hour at 37°C before aspirating excess solution. Cells were cultured in Essential 8 (E8) medium, prepared in-house according to the original formulation described by Chen et al. (2011) supplemented with 100 ng/mL bFGF. After initial thaw and plating, cells were expanded for 3 days, then dissociated with TrypLE (Gibco), replated on freshly coated Matrigel plates, and grown for a further 4 days in E8 supplemented with 30 ng/mL bFGF before use in differentiation.

All cell lines were authenticated by providence, fluorescent protein expression, and confirmed to be karyotypically normal and mycoplasma free within two passages of experimental use.

HES2 hESCs (Reubinoff et al., 2000) (HES-2 (RRID:CVCL_D093)) were previously targeted at the human ROSA locus to constitutively express tdRFP (Irion et al. 2007). HES2 hESCs expressing humanized GFP from the AAVS1 locus were previously generated by AAV2 mediated integration (Yang et al., 2008). PGPC17-11 iPSCs (karyotype: 46XY) (Hildebrandt et al., 2019) were obtained from Dr. James Ellis (SickKids Research Institute, Toronto, ON, Canada). ESI-17 (BioTime, Alameda, CA, USA) (Chen et al., 2015) was obtained from Dr. Michael LaFlamme. CHOPWT17 (RRID: CVCL_C9JD) were obtained from Dr. Deborah French (Maguire, J.A. et al., 2016).

The hPSC studies were subject to approval by the Stem Cell Oversight Committee (Canadian Institutes of Health Research).

#### Animals

All animal experiments were conducted under protocols approved by the Animal Care Committee of the University Health Network (Toronto, Canada) and the University of Western Ontario, and were carried out in accordance with the guidelines set by the Canadian Council on Animal Care (CCAC). NOD-SCID gamma (NSG; NOD.Cg-*Prkdc*scid*Il2rg*tm1Wjl/SzJ) mice (The Jackson Laboratory) aged 6–10 weeks were used as recipients for all transplantation experiments. Mice were housed in a specific pathogen-free (SPF) facility under a 12-hour light/dark cycle with ad libitum access to water and irradiated chow. Male and female mice were used in equal proportions. Animals were randomized to experimental groups. Sample sizes for each experiment are described in the Figure Legends. For hindlimb ischemia studies, male mice were randomly assigned to treatment groups. Investigators were blinded to group allocation during outcome assessment.

## Key Resources Table

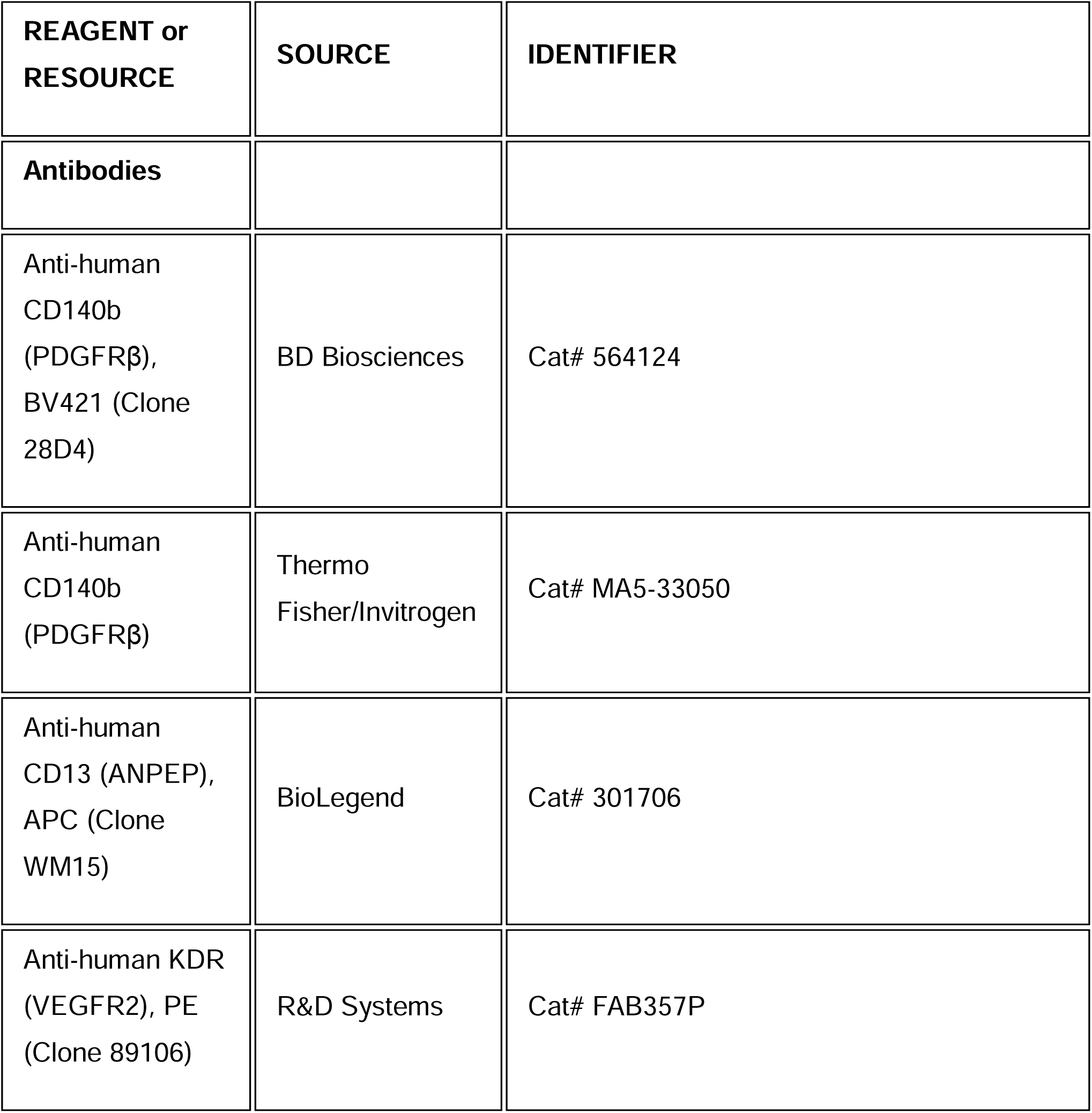
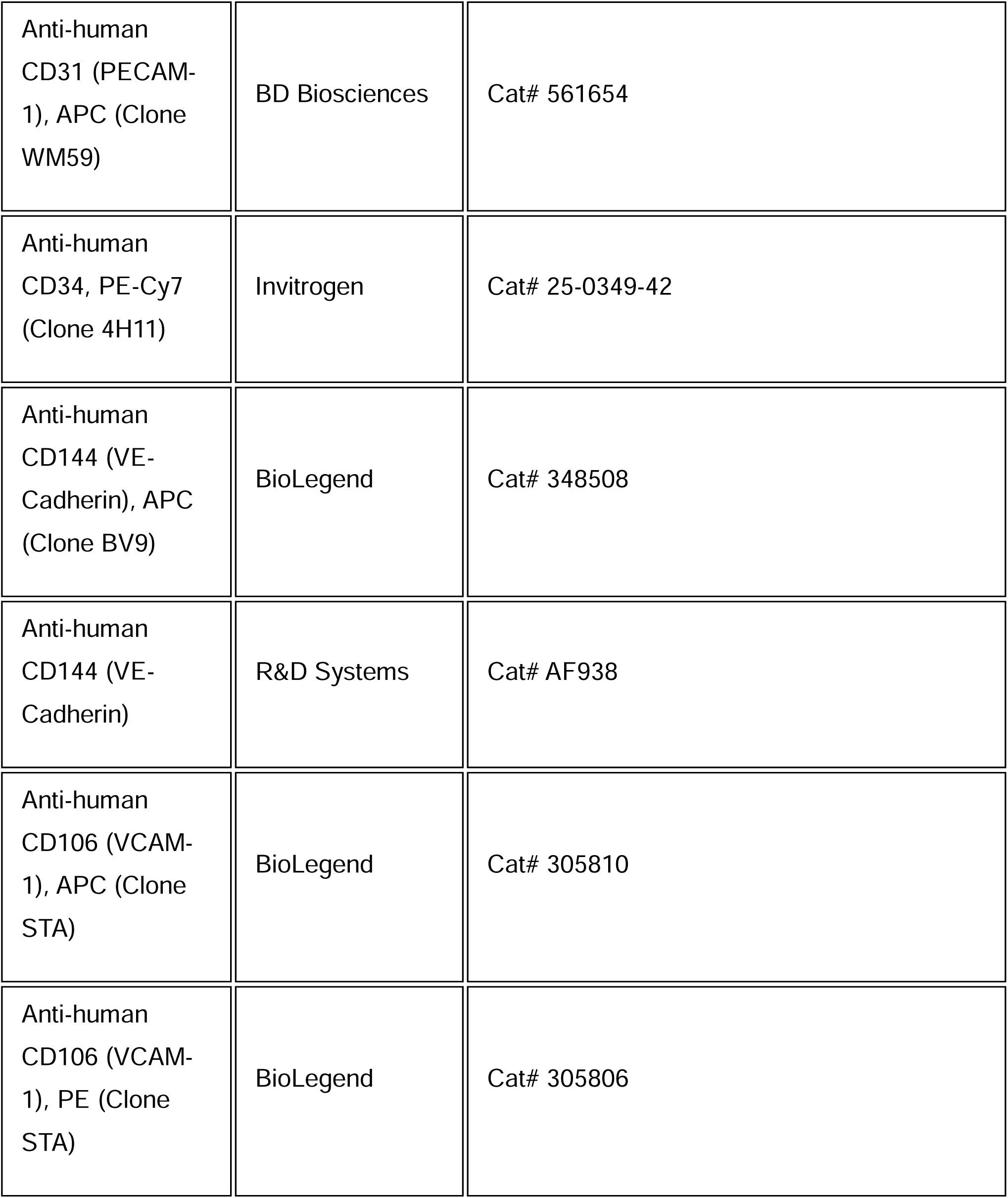
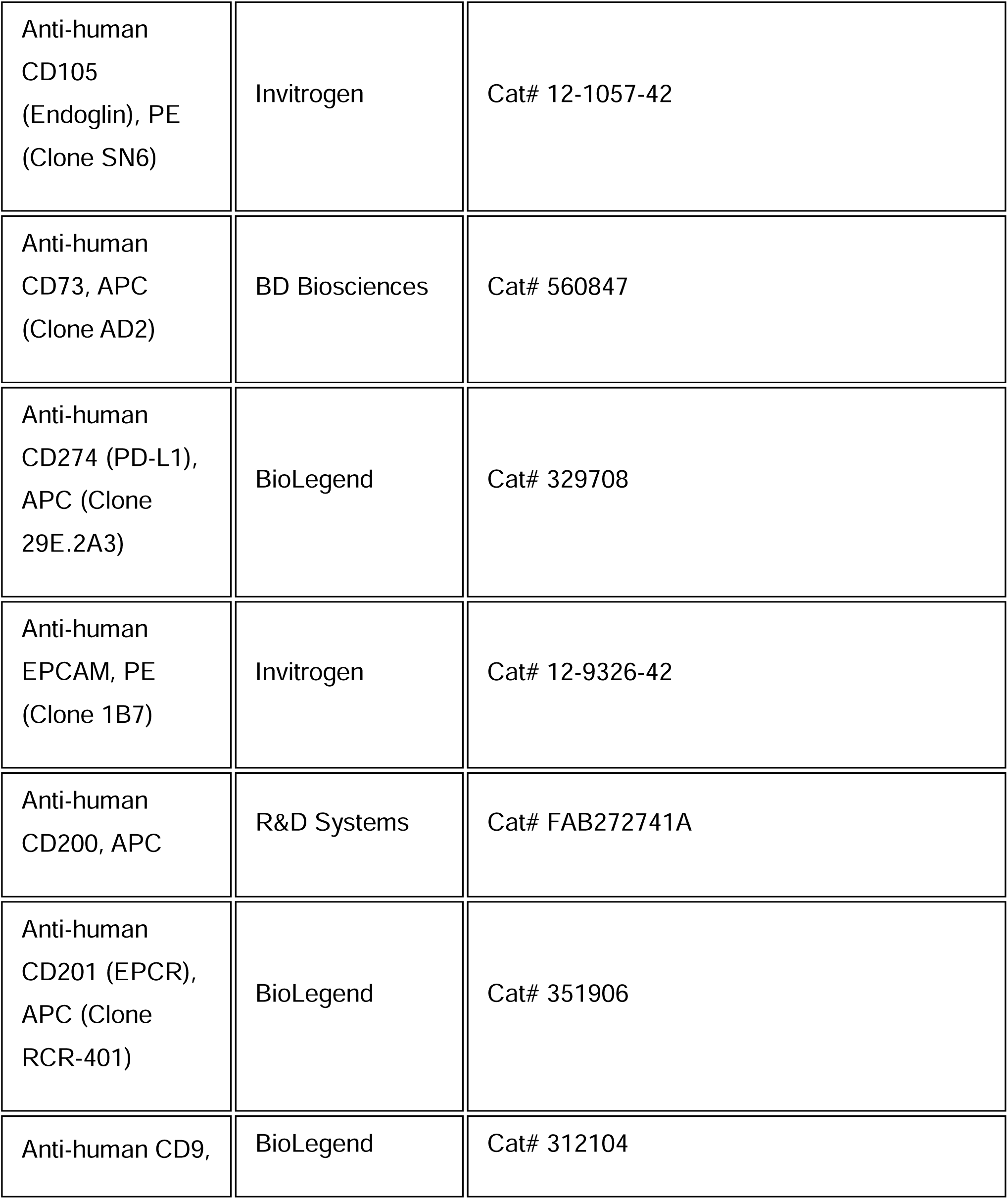
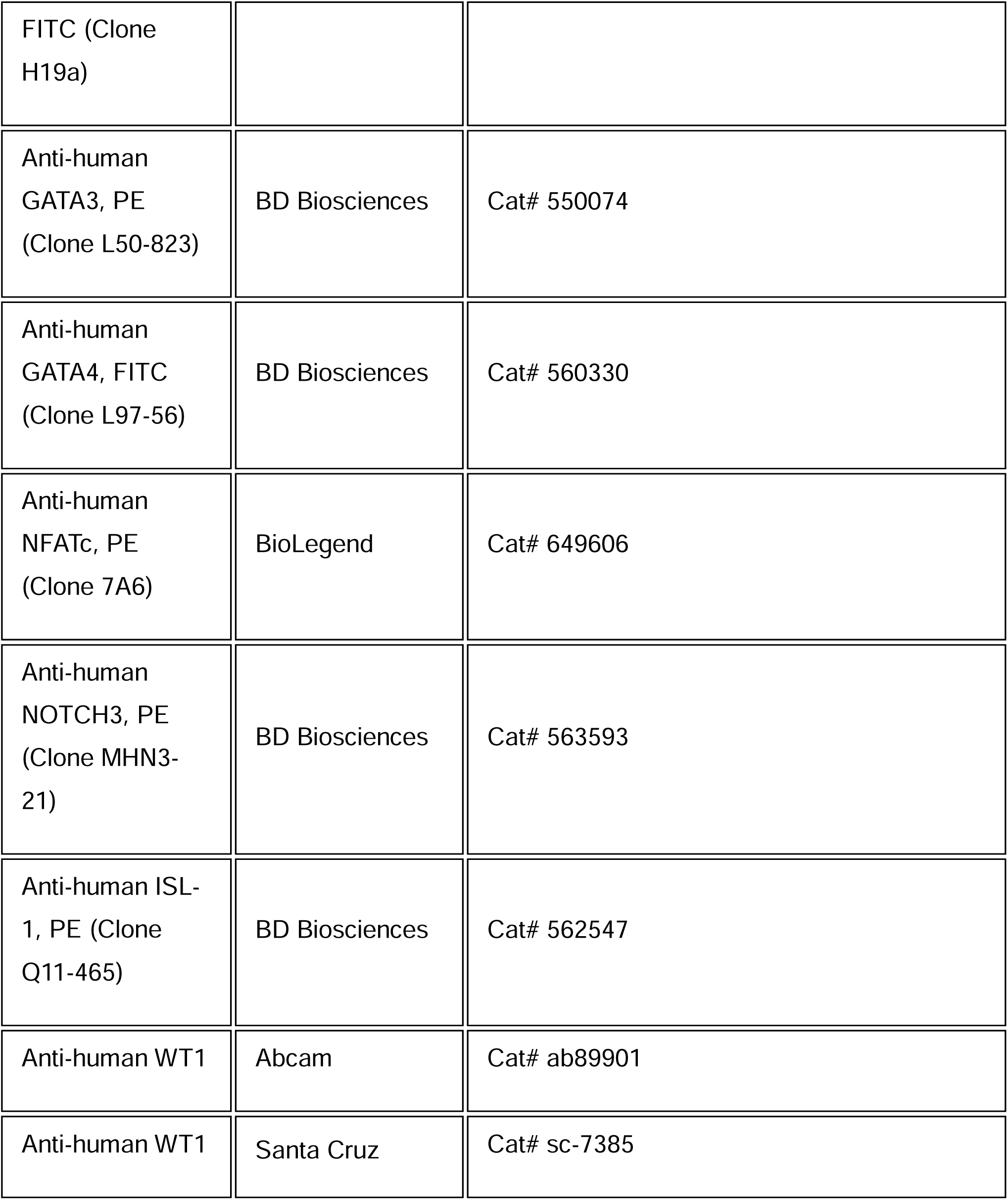
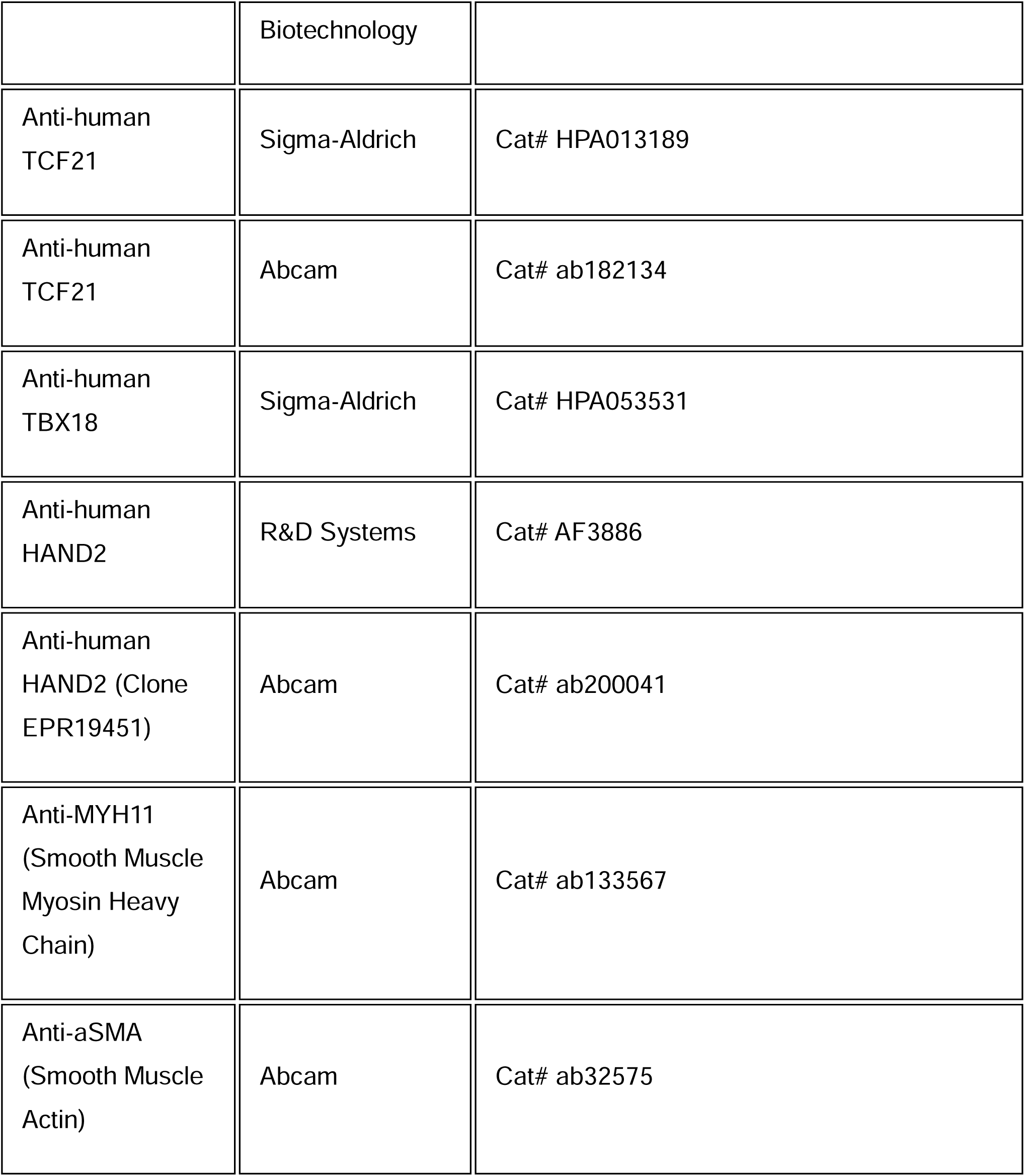
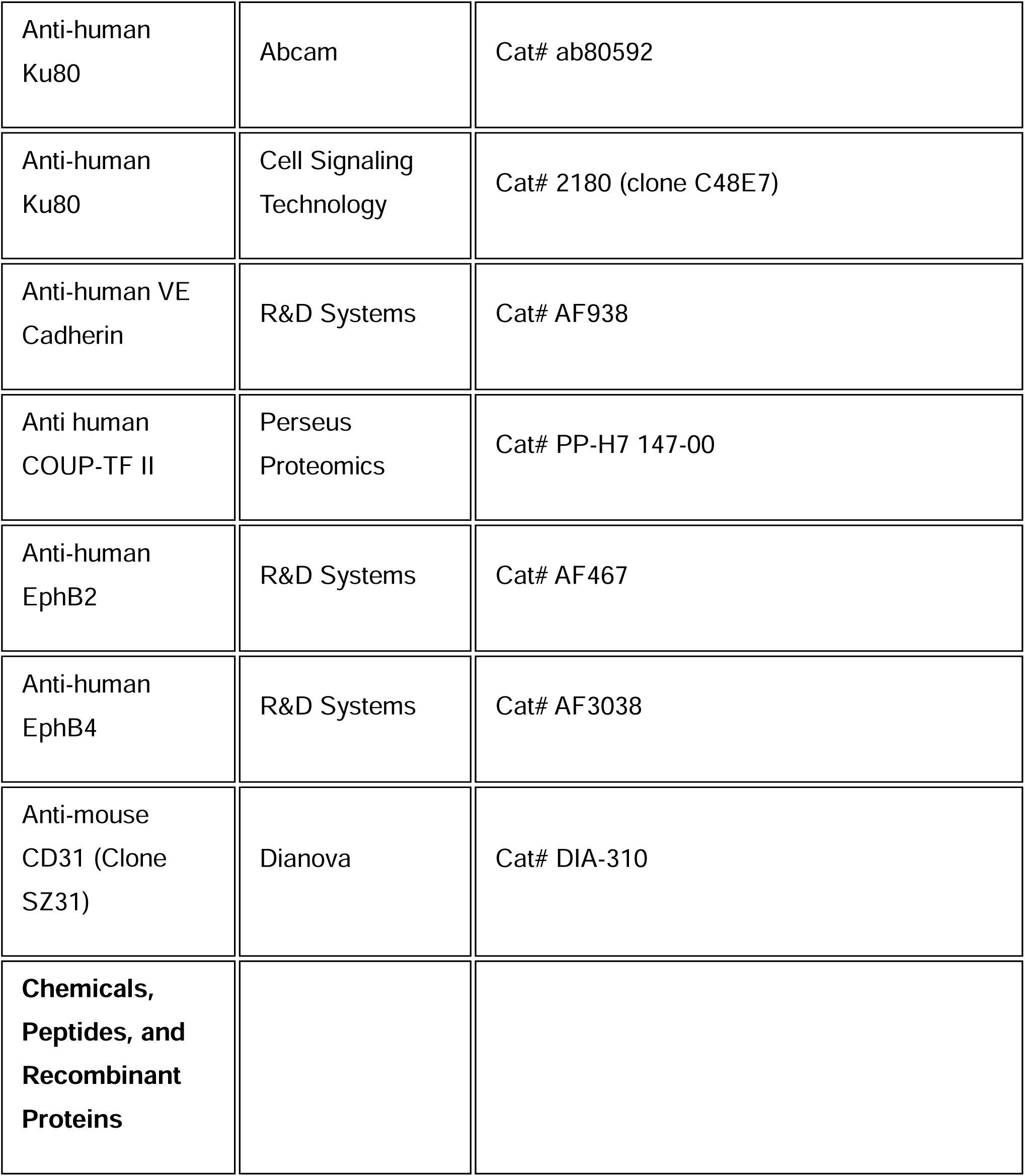
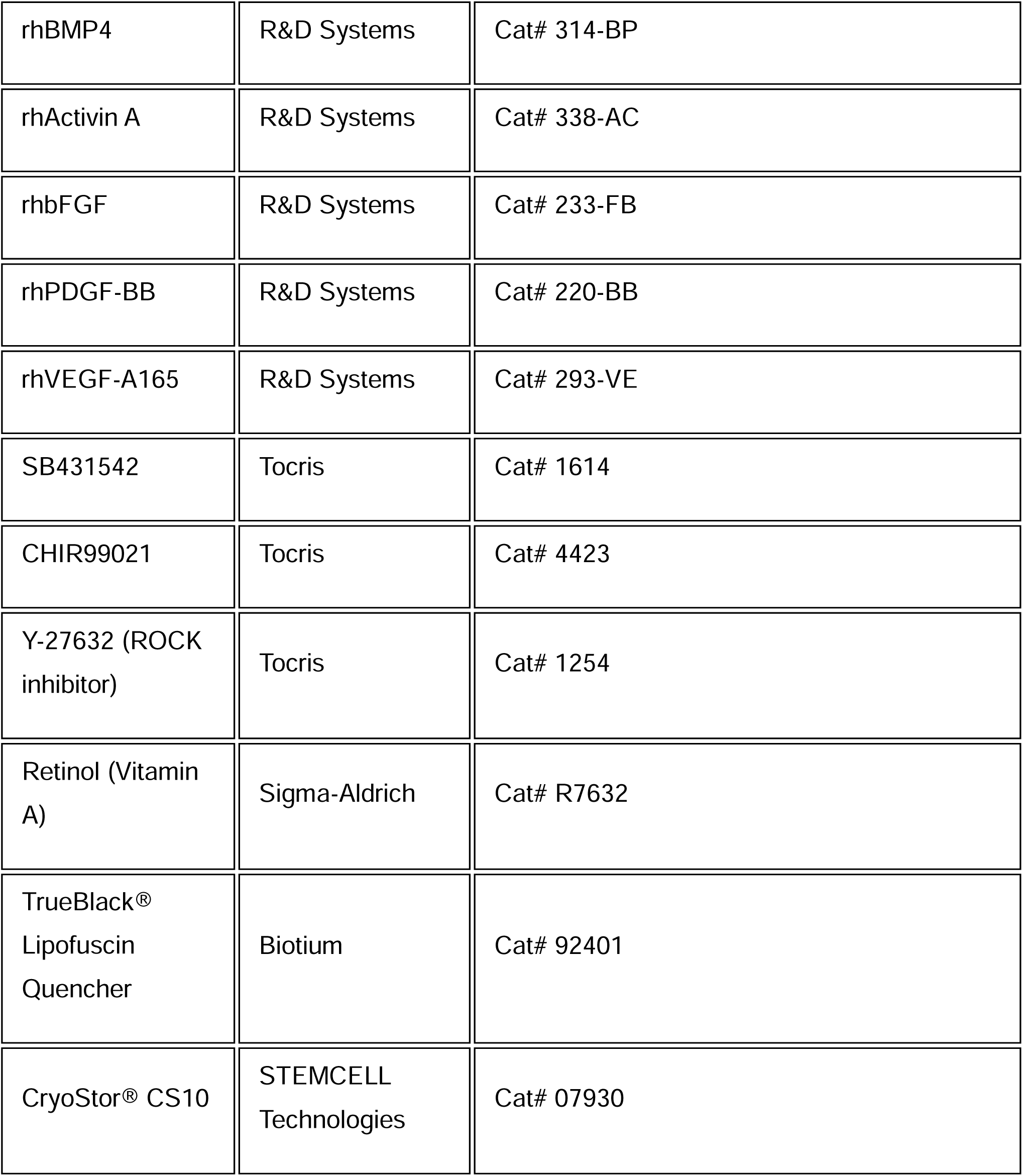
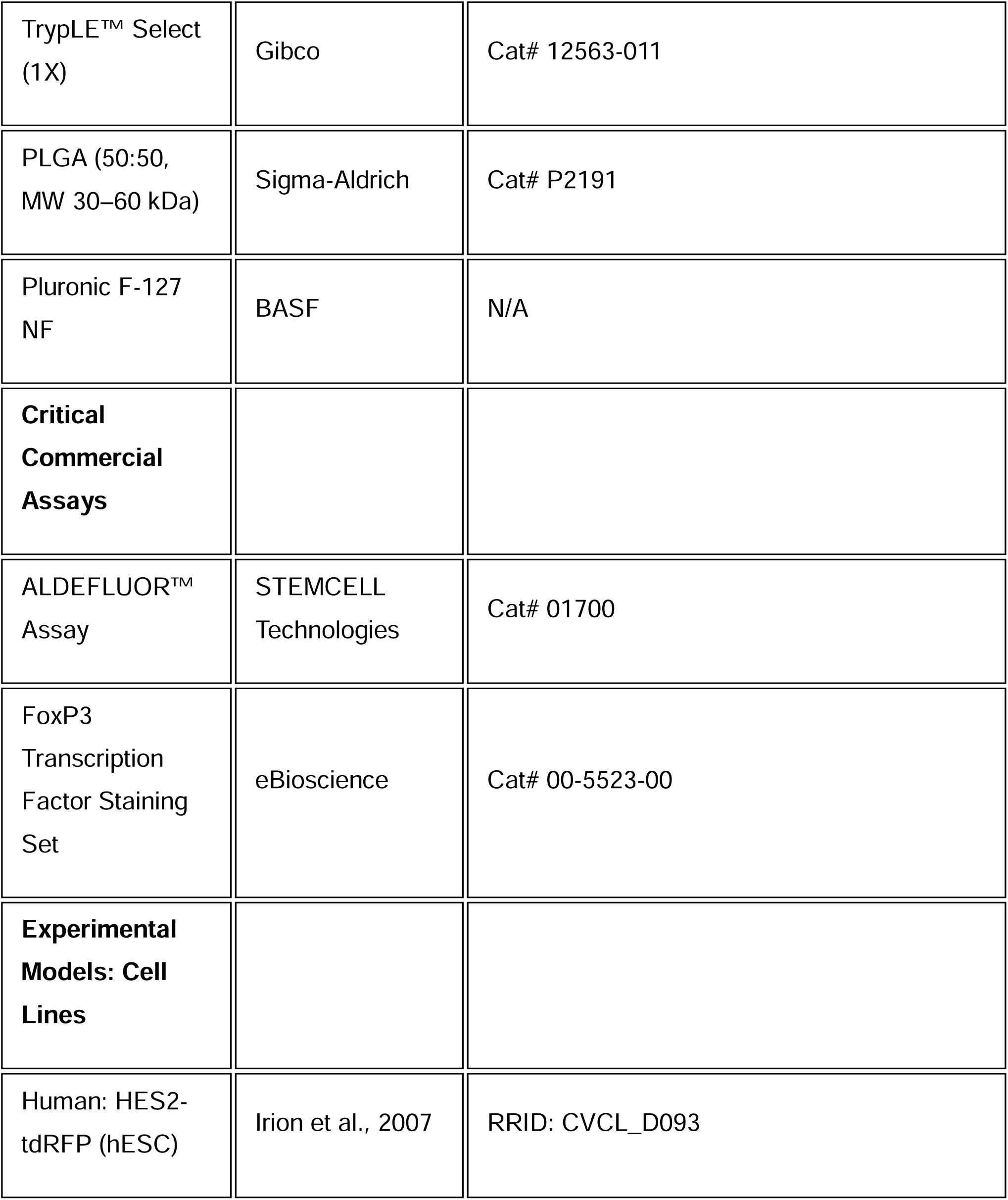
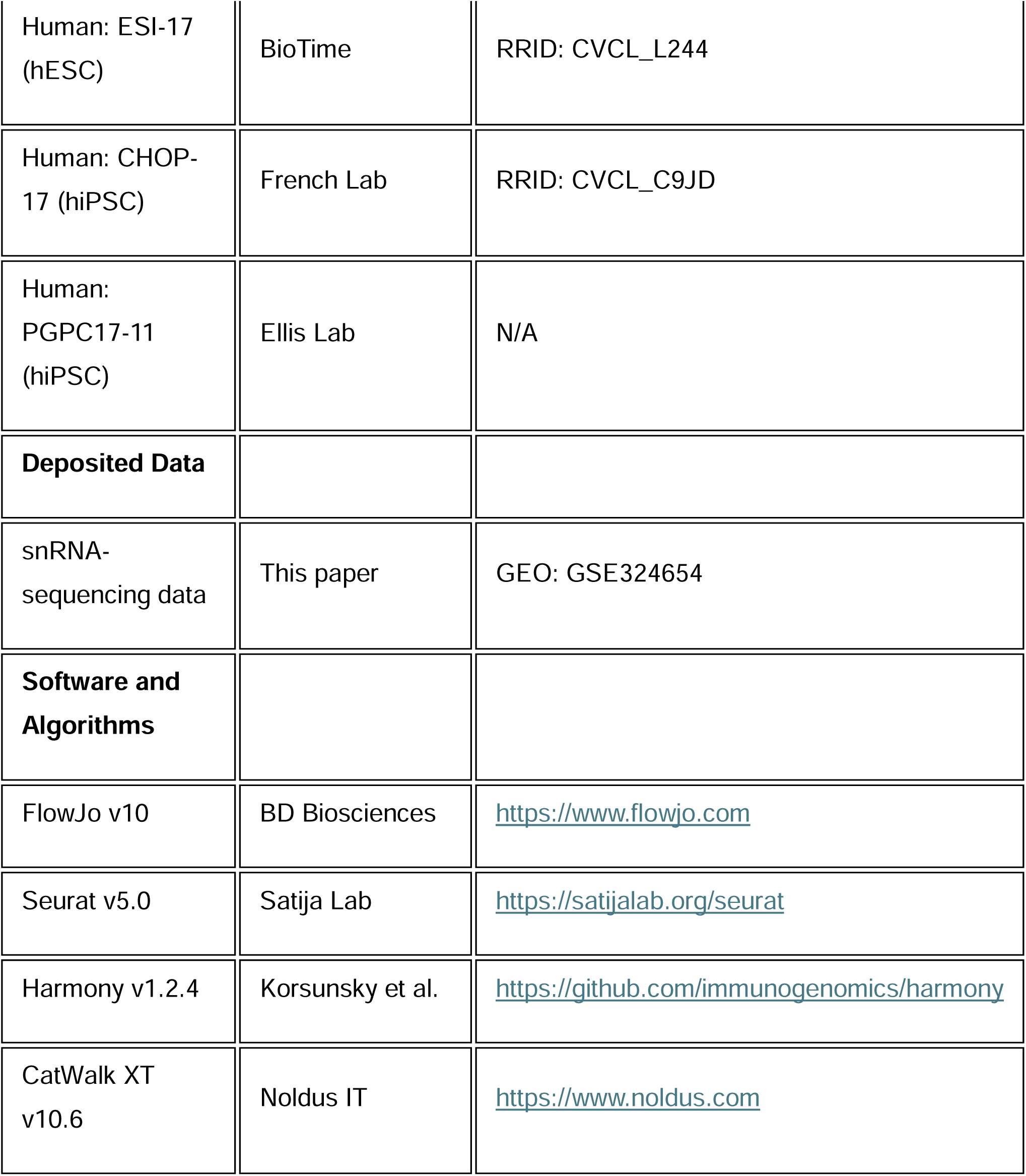
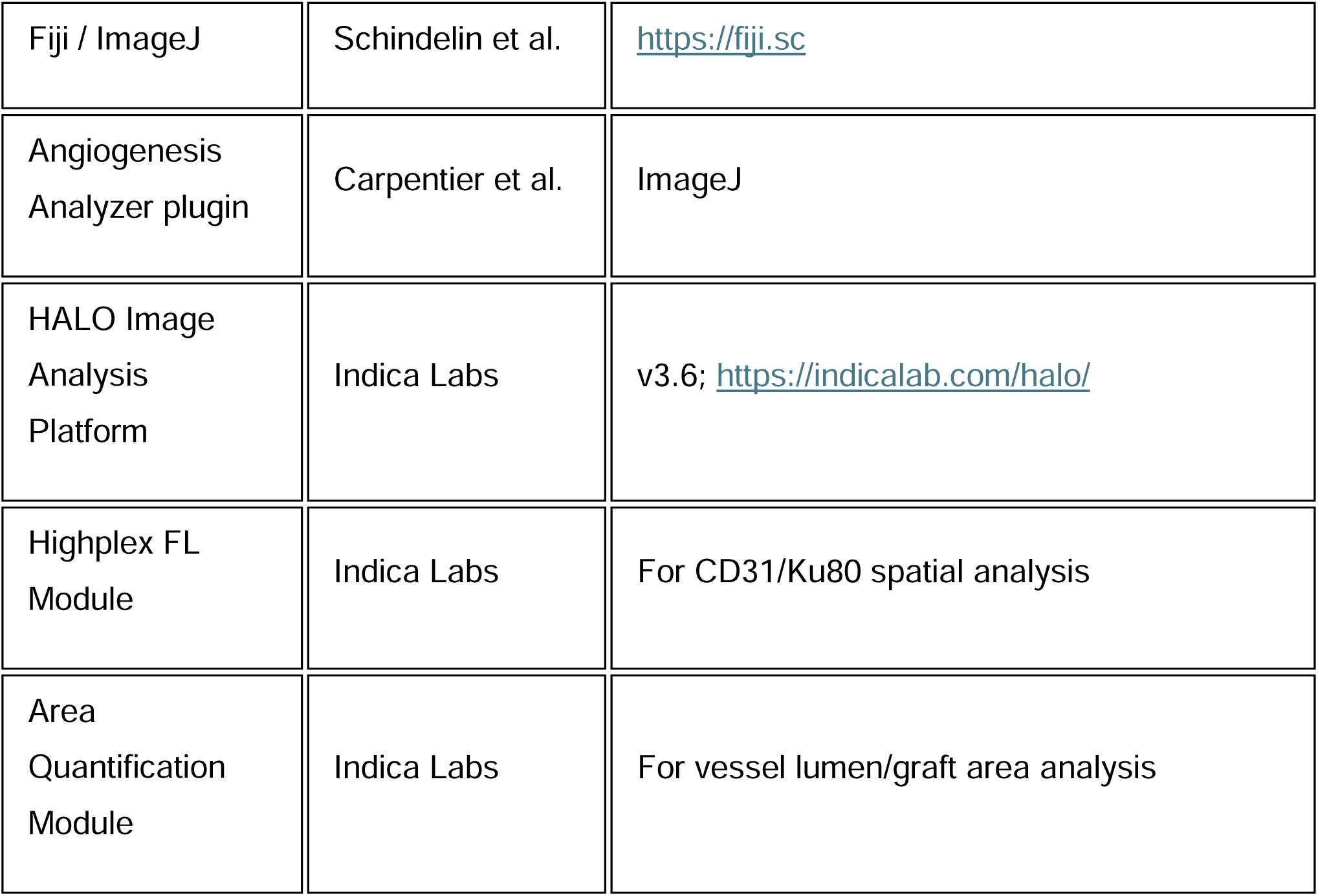

## METHOD DETAILS

### hPSC Differentiation: VP Cell Generation

Vascular progenitor (VP) cells were generated using a modified version of our previously published epicardial/cardiac fibroblast differentiation protocol (Fernandes et al., 2023). The protocol proceeds through the following staged inductions over 23 days under low oxygen conditions (5% O_2_, 5% CO_2_) unless otherwise specified. The base differentiation medium throughout consists of 25% StemPro-34 / 75% IMDM (Thermo Fisher Scientific) supplemented with 2 mM L-glutamine, 50 μg/mL ascorbic acid (Sigma), 150 μg/mL transferrin (Roche), and 400 μM monothioglycerol (MTG; Sigma), hereafter referred to as Basal Medium.

**Stage 0 — Embryoid Body (EB) Formation, Days 0–1:** hPSCs were dissociated into single cells by incubation with TrypLE (Thermo Fisher Scientific) for 2–3 minutes at 37°C until MEF feeders visibly detached. hPSC clusters were further dissociated by trituration and washed twice in wash medium (IMDM + 0.5%w/v BSA). Cells were filtered through a 40 μm cell strainer (Corning) and collected by centrifugation at 1,000 rpm for 5 minutes. Single cells were then aggregated to form EBs by culture in Basal Medium supplemented with 5 μM Y-27632 ROCK inhibitor (Tocris) and 1 ng/mL rhBMP4 (R&D Systems) at 2 × 10^6^ cells/mL in 4 mL per 6 cm petri-grade dish (non-tissue culture treated) on an orbital shaker (MaxQ 2000; Thermo Fisher Scientific) at 70 rpm. Cultures were maintained at low oxygen (5% O_2_, 5% CO_2_) for 18 hours.

**Stage 1 — Mesoderm Specification, Days 1–4:** On day 1, EBs were harvested through a 100 μm strainer and pelleted at 600 rpm for 5 minutes. EBs were transferred to Basal Medium supplemented with 5 ng/mL bFGF (R&D Systems), and rhBMP4 and rhActivin A at concentrations optimized per cell line (for HES2: 3 ng/mL BMP4 and 1 ng/mL Activin A to generate second heart field [SHF] mesoderm). EBs were plated on polyHEME-coated 6-well plates in static low-oxygen culture. On day 4, EBs were dissociated using TrypLE for 5 minutes at 37°C, washed with wash medium containing DNase, and cells counted. Mesoderm quality was assessed by flow cytometry for CD140a (PDGFRα) and ALDH activity (ALDEFLUOR assay; Stem Cell Technologies) and CD235a/b (glycophorin A/B). SHF mesoderm, optimal for pro-epicardial induction, is defined by 30% CD140a^+^ALDH^+^ cells with <15% CD140a^−^ cells.

**Stage 2 — Pro-Epicardial Specification, Days 4–8:** Day 4 cells were replated as a monolayer on 0.1% gelatin-coated tissue culture plates at 6 × 10^5^–8 × 10^5^ cells/mL in Basal Medium containing 10 ng/mL BMP4, 1 μM CHIR99021, 2 μM retinol (vitamin A; Sigma), and 6 μM SB431542 (ALK4/5/7 inhibitor; Tocris). On day 6, medium was fully replaced with Basal Medium supplemented with 6 μM SB431542 only. Medium was changed every 48 hours.

**Stage 2.1 — Epicardial Development, Days 8–12:** On day 8, the monolayer was dissociated by incubation with Collagenase B (1 mg/mL in IMDM; Roche) for 1 hour at 37°C, which typically caused the monolayer to detach in sheets. Cells were collected at 400 rpm for 5 minutes, then further dissociated enzymatically with TrypLE for 5 minutes at 37°C, stopped by 10-fold dilution in wash medium, followed by a gentle trituration/vortex. Cells were replated at 3 × 10^5^ cells/mL on gelatin-coated plates in Basal Medium with 6 μM SB431542. Cultures were maintained in low oxygen through day 12.

**Stage 2.2 — Epicardial Expansion, Days 12–17:** On day 12, cells were again dissociated using Collagenase B (1 hour at 37°C) followed by TrypLE (5 minutes), washed, and replated at 1.5 × 10^5^ cells/mL on gelatin-coated plates in Basal Medium with 6 μM SB431542. Medium was changed every 48 hours. By day 17, cells display a mature epicardial morphology with cobblestone appearance and express WT1, TBX18, and TCF21.

**Stage 3 — Vascular Progenitor Induction, Days 17–23:** On day 17, epicardial cells were harvested by Collagenase B treatment (1 hour, 37°C) followed by TrypLE dissociation (5 minutes). Cells were replated on gelatin-coated dishes at 3 × 10^5^ cells/mL in Basal Medium supplemented with 10 ng/mL bFGF, 10 ng/mL PDGF-BB (R&D Systems), 1 ng/mL TGFβ1 (R&D Systems), and 5 ng/mL EGF (R&D Systems) for 72 hours (days 17–20; EMT induction step). On day 20, medium was fully aspirated from the newly formed adherent monolayer and replaced with Basal Medium containing 10 ng/mL bFGF, 10 ng/mL PDGF-BB, and 6 μM SB431542 for an additional 72 hours (days 20–23; differentiation/expansion step). Cultures were maintained under low oxygen conditions throughout Stage 3. At day 23, cells were dissociated with TrypLE (5 minutes at 37°C) and either used immediately for downstream analysis or cryopreserved.

**Fibroblast Control Differentiation:** For parallel generation of cardiac fibroblast-like cells (CFs) used as comparators, the Stage 3 induction step from days 17–20 used: 30 ng/mL bFGF, 6 μM SB431542, 1 μM CHIR99021, and 500 nM retinoic acid (RA; Sigma), as previously described (Fernandes et al., 2023).

### VP Cell Cryopreservation and Thawing

Day 23 VP cells were cryopreserved in CryoStor CS10 (Stem Cell Technologies) at 1–10 × 10^6^ cells/mL in cryovials and stored in vapor-phase liquid nitrogen. For thawing, vials were rapidly thawed at 37°C, and cells were diluted dropwise into pre-warmed IMDM + 0.5% BSA, centrifuged at 1,200 rpm for 5 minutes, and resuspended in VP culture medium (Basal Medium supplemented with 10 ng/mL bFGF and 10 ng/mL PDGF-BB) on 1% Matrigel-coated plates. Cells were allowed to recover for 48 hours before use. Quality control was assessed at day 1–3 post-thaw by flow cytometry for CD140b, CD13, and KDR (minimum QC thresholds: ≥60% CD140b^+^CD13^+^, ≥20% KDR^+^).

### VEGF-Induced Endothelial Differentiation

For VEGF-induced endothelial differentiation, VP cells (day 23) and earlier-stage populations were cultured in EGM-2 (Lonza) supplemented with 100 ng/mL recombinant human VEGF-A (R&D Systems) for 48 hours, then analyzed by flow cytometry for CD31 and CD34.

For Matrigel tube formation assays, cells were labeled with Calcein AM (Thermo Fisher; 1 μg/mL, 30 minutes at 37°C), then seeded at 1.5 × 10^4^ cells/well onto growth factor–reduced Matrigel (Corning; 50 μL/well in a 96-well plate) and imaged after 48 hours using fluorescence microscopy. Network complexity was quantified using the Angiogenesis Analyzer plugin in Fiji (ImageJ) for junctions, segments, branches, and meshes.

### Calcium Imaging

VP cells and cardiac fibroblasts cells were loaded with Fluo-4 AM calcium indicator (Thermo Fisher; 5 μM, 30 minutes at 37°C), washed, and treated with 500 nM endothelin-1 (ET-1; Sigma). Rapid calcium flux was measured by live fluorescence imaging on an inverted widefield microscope.

### Flow Cytometry and Cell Surface Marker Analysis

For surface marker analysis, cells were dissociated with TrypLE, washed in PBS + 2% FBS (FACS buffer), and incubated with fluorophore-conjugated antibodies for 20 minutes at 4°C in the dark. For intracellular transcription factor staining, cells were first surface stained, then fixed and permeabilized using the FoxP3 Transcription Factor Staining Buffer Set (eBioscience/Thermo Fisher) according to the manufacturer’s instructions, followed by intracellular antibody incubation for 45 minutes. DAPI was used for live/dead cell gating. Samples were analyzed on an LSRFortessa X-20 (BD Biosciences) or sorted on a FACSAria III (BD Biosciences). Data were analyzed using FlowJo software (v10; BD Biosciences).

Antibodies used for flow cytometry can be found in the key resource table. ALDH1A2/RALDH2 enzyme activity was monitored using the ALDEFLUOR assay (STEMCELL Technologies), performed according to the manufacturer’s instructions. All antibodies were validated using appropriate isotype or fluorescence-minus-one (FMO) controls. Specific catalog numbers and vendors are listed in the Key Resources Table.

### PLGA Nanoparticle Fabrication and Loading

Poly(lactic-co-glycolic acid) (PLGA) nanoparticles loaded with recombinant human VEGF-A_165_ or BSA (control) were fabricated using a water/oil/water double emulsion solvent evaporation method, as previously described (Shoichet lab, University of Toronto). Briefly, PLGA (50:50, MW 30–60 kDa; Sigma) and 0.05 wt % Pluronic F-127 NF (BASF) were dissolved in dichloromethane and emulsified with an aqueous VEGF-A solution. The emulsion was then transferred to an aqueous polyvinyl alcohol (PVA) solution and homogenized to form the double emulsion. Solvent was evaporated by stirring gently overnight. Nanoparticles were washed by centrifugation, lyophilized, and stored at −80°C until use. Nanoparticle size (200–3400 nm) and polydispersity index (PDI < 0.1) were characterized by dynamic light scattering. VEGF encapsulation efficiency (249 ng/mg PLGA) was determined by ELISA after dissolving the nanoparticles in chloroform/DI water to release the encapsulated VEGF. Prior to transplantation, lyophilized nanoparticles were reconstituted and mixed with VP cell suspensions in Matrigel.

### Mammary Fat Pad Transplantation

NSG recipient mice were anesthetized with isoflurane (2–3%). Mammary glands were localized without incision by externally marking nipples followed by wiping the ventral surface with 70% isopropanol. Cell and Matrigel (final concentration 5-7.5mg/ml) suspensions were prepared on ice as 50–100 μL of the cell-Matrigel suspension (containing 1×10 –10×10 VP cells, or Stage 1/2 cells as specified per experiment). Cell and Matrigel suspensions, loaded in an ice cold 26G1/2 insulin syringe were injected directly into the fat pad taking care to remain on the anatomical mammary line. Up to four grafts were placed per recipient specifically left and right thoracic (between nipple 2 and 3, adjacent to mammary artery) and left and right inguinal (between nipple 4 and 5). Grafts were analyzed at days 3, 7, 14, 26, and 28 post-transplantation by histology as described below. Dose-response experiments compared 1×10, 5×10, and 10×10 cells per transplant; 5×10 was used as the standard dose for subsequent experiments. For VEGF nanoparticle co-delivery experiments, PLGA nanoparticles loaded with either VEGF or BSA control were mixed with the cell-Matrigel suspension immediately prior to injection. All animal procedures were performed in accordance with institutional guidelines and approved by the animal care committee.

### Skeletal Muscle Transplantation (Uninjured Hindlimb)

For transplantation into uninjured skeletal muscle, NSG mice were anesthetized with isoflurane. A small incision was made over the medial aspect of the hindlimb, and 5 × 10^6^ VP cells in 50 μL of Matrigel were injected directly into the vastus intermedialis using a 26-gauge needle. Grafts were analyzed at 2 weeks, 4 weeks, and 12 weeks post-transplantation. At each endpoint, mice were euthanized, and the gastrocnemius muscle was dissected for histological analysis.

### Hindlimb Ischemia Models

#### Mild ischemia model (femoral artery ligation)

Mild hindlimb ischemia was induced in NSG mice by permanent ligation of the femoral artery just distal to the profunda femoris using a 7-0 Prolene suture, without resection. This procedure induces moderate ischemic injury primarily to the vastus intermedialis muscle. VP cells (5 × 10^6^) were transplanted into the ischemic vastus intermedialis 7 days following ligation. Successful ischemia induction was confirmed at harvest by Masson’s trichrome staining showing increased fibrosis in ligation-injured compared to control (uninjured contralateral) muscle. Grafts were analyzed at 4 weeks post-transplantation. For co-transplantation with nanoparticles, VP cells (1 × 10^6^) were mixed with VEGF-A-loaded or BSA-loaded PLGA nanoparticles (NPs) prior to injection.

#### Severe ischemia model (femoral artery resection)

Severe hindlimb ischemia was induced by complete surgical resection of the femoral artery, from the inguinal ligament to the popliteal bifurcation. (Arpino et al. 2021).This procedure results in extensive ischemic injury to the lower limb musculature. Within 1 week of injury, 5 × 10^6^ VP cells (BSA-NPs or in combination with VEGF-A-loaded nanoparticles) in Matrigel were injected into the medial gastrocnemius and soleus of the injured limb. Matrigel alone (without cells) was injected as a control. The 1-month and 3-month cohorts were analyzed for blood flow perfusion, functional gait outcomes, histological analysis of vasculature, and myofiber morphology.

### Laser Speckle Perfusion Imaging

Blood flow perfusion in the hindlimb was monitored longitudinally using a laser speckle perfusion imaging system (Moor Instruments) (Li et al. 2020). Mice were anesthetized with isoflurane and placed in a supine position on a warming pad at 37°C. Plantar surfaces of both hindpaws were imaged prior to surgery (baseline) and at 1, 2, 3, and 4 weeks and 1-, 2-, and 3-months post-transplantation. Mean flux values were measured in standardized regions of interest (ROIs) encompassing the ischemic and contralateral paws. Blood flow is expressed as the ratio of ischemic-to-contralateral limb flux to normalize for inter-animal variability and body temperature effects.

### Gait Analysis (CatWalk XT)

Gait and weight-bearing were assessed using the CatWalk XT automated gait analysis system (Noldus Information Technology) at 1- and 3-months post-transplantation Citation: Said et al. Adv Health Care Mater 2019, PMID 30785239. Mice were trained to cross the CatWalk walkway for three runs prior to baseline recording. Animals were required to complete a minimum of three valid runs (uninterrupted crossings within 10 seconds). The following parameters were analyzed for the ischemic right hindpaw compared to the contralateral left hindpaw: print area, maximum contact area, duty cycle, maximum intensity, and maximum contact maximum intensity. Data were analyzed using the CatWalk XT software (v10.6). Statistical comparisons across treatment groups were performed using a generalized linear mixed model (GLMM) with individual mouse as a random effect. Reported n values reflect total runs across all animals per group.

### Immunohistochemistry and Histology

Explanted tissues (fat pad, gastrocnemius, soleus, vastus intermedialis) were fixed in 10% neutral buffered formalin or 4% PFA for fat pad explants for 24 hours at room temperature, processed through graded ethanol and xylene series, and embedded in paraffin. Sections of 5 μm thickness were cut using a rotary microtome and mounted on SuperFrost Plus slides (Thermo Fisher).

For immunofluorescence, sections were dewaxed, rehydrated, and subjected to heat-induced epitope retrieval (HIER) in citrate buffer (pH 6.0; 20 minutes at 95°C) or Tris-EDTA buffer (pH 9.0; 20 minutes at 95°C) depending on the primary antibody. Sections were blocked with 5% normal donkey serum in PBS + 0.1% Triton X-100 for 1 hour at room temperature, then incubated with primary antibodies overnight at 4°C. The following primary antibodies were used for histology: anti-human CD31 (clone JC70A, DAKO;), anti-human Ku80/XRCC5 (rabbit polyclonal, Abcam, ab80592; 1:200) for human-specific nuclear identification, anti-mouse CD31 (rat monoclonal, Dianova; 1:20), anti-MYH11 (rabbit polyclonal, Abcam; 1:200), anti-CD144/VE-cadherin (rabbit anti-human, Abcam; 1:100), anti-CD140b/PDGFRβ (rabbit anti-human; 1:100), anti-laminin (rabbit polyclonal, MilliporeSigma; 1:30), anti-RFP (for tdRFP tracking; rabbit polyclonal, 1:200), anti-GFP (for eGFP tracking). Alexa Fluor-, horseradish peroxidase-, or alkaline phosphatase-conjugated secondary antibodies (donkey anti-rabbit, anti-mouse, anti-rat; 1:500; Thermo Fisher) were used. Sections were counterstained with DAPI (1 μg/mL) and mounted with ProLong Gold (Thermo Fisher). Images were acquired on a Zeiss LSM 800 confocal microscope or Leica Stelaris 1 confocal microscope.

For Masson’s trichrome staining, deparaffinized sections were stained according to standard protocol (Sigma Aldrich Trichrome Stain Kit) to assess fibrosis (blue), muscle (red), and nuclei (dark brown/black). Myofiber cross-sectional area and fibrotic area were quantified from trichrome images using Fiji (ImageJ) with the Feret’s diameter measurement or particle analysis tools.

Quantitative image analysis was performed in Fiji (ImageJ). For CD31/Ku80 dual staining, the percentage of CD31^+^ cells among total Ku80^+^ human cells was computed per section by automated thresholding of each channel. Human CD31+Ku80+ vascular area (mm²) and vessel lumen density (lumen/mm²) were quantified across the total graft area using automated particle detection.

### Digital Image Analysis and Histological Quantification

Whole-slide images of transplanted tissues were acquired using an Aperio slide scanner and then analyzed using the HALO image analysis platform (Indica Labs, v3.6). Vessel lumen density and total graft area (mm^2^) were measured using the Area Quantification module. For CD31 and Ku80 dual-staining, the Highplex FL module was employed to quantify the percentage of human (Ku80^+^) cells that co-expressed the endothelial marker CD31.

### RT-qPCR

RNA was isolated from bulk population from *in vitro* hPSC-derived samples using a RNAqueous-micro kit (Invitrogen). After RNA isolation the eluate of each column, containing up to 1 μg of RNA, was treated with DNAse to remove genomic DNA followed by cDNA synthesis (iSCRIPT, BioRad). Reverse-transcription quantitative PCR (RT-qPCR) was performed using a CFX384 Touch Real-Time instrument (BioRad), SsoAdvanced Universal SYBR Green Supermix (BioRad, Cat#204057) with primers described in the key resource table, following manufacturer instructions. Gene expression relative to TBP was determined from technical duplicates, relative copy number and reaction efficiency. Genomic DNA standards were made in lab from wild type HES2 hESC cells and used in a 10-fold dilution series ranging from 2.5 pg/μL to 25 ng/μL. Bar graphs of gene expression were generated in Prism version 10 (Graphpad).

### Single-Cell RNA Sequencing

Single-nucleus RNA sequencing (snRNA-seq) was performed on day 23 VP cells prior to transplantation and on cells isolated from fat pad and hindlimb grafts at 1-month post-transplantation. For graft cell isolation, freshly explanted tissues were minced and dissociated using Collagenase IV (1 mg/mL in IMDM; Thermo Fisher) for 45 minutes at 37°C with intermittent agitation, followed by filtration through a 70 μm strainer.

Single nucleus libraries were prepared using the 10x Genomics Chromium Single Cell 3’ kit (v3.1) according to the manufacturer’s instructions, targeting 5,000–10,000 cells per sample. Sequencing was performed on an Illumina NextSeq 2000 or NovaSeq 6000 to a depth of ≥30,000 reads per cell. Raw sequencing reads were processed using Cell Ranger (10x Genomics, v7.0) with alignment to a human (GRCh38) reference genome. Ambient RNA removal was performed using SoupX (v1.6). Doublet detection was performed using DoubletFinder (v2.0).

Downstream analysis was performed in R (v4.3) using Seurat (v5.0). Cells were filtered based on the following quality metrics: ≥200 and ≤6,000 genes detected, <20% mitochondrial gene content for dissociated cells, and <5% for nuclear preparations. Data from individual samples were normalized using SCTransform (v2), and integration was performed using Harmony (v1.2.4) to correct for batch effects. Dimensionality reduction was performed by principal component analysis (PCA) followed by UMAP embedding. Unsupervised clustering was carried out at resolution 0.5 using the Leiden algorithm. Cell type annotation was based on the expression of canonical marker genes: endothelial cells (*PECAM1, CDH5, FLI1*), pericytes/smooth muscle (*PDGFRB, ACTA2, RGS5, MYH11*), mesenchymal (*VIM, DCN*), and transitional populations (RALDH2). Differential gene expression between clusters or conditions was assessed using the Wilcoxon rank-sum test with Bonferroni correction. Gene set enrichment analysis (GSEA) was performed using the fGSEA package against MSigDB hallmark and curated vascular gene sets. Cell-type composition differences between fat pad and hindlimb-derived endothelial cells were assessed after sub-clustering of CD31^+^ endothelial populations at resolution 0.3, yielding 13 distinct endothelial subclusters that were annotated as arterial, venous, capillary, tip cell, proliferating, lymphatic, or mixed subtypes based on established gene signatures.

### QUANTIFICATION AND STATISTICAL ANALYSIS

All data are presented as mean ± SEM unless otherwise stated. Statistical analyses were performed in GraphPad Prism (v10) or R (v4.4.1). For comparisons between two groups, unpaired two-tailed Student’s t-test was used. For comparisons among three or more groups, one-way ANOVA with Tukey’s honest significant difference (HSD) post-hoc test was applied. Longitudinal laser Doppler data were analyzed by two-way ANOVA with Šídák’s multiple comparisons correction. CatWalk gait data were analyzed using a generalized linear mixed model (GLMM) with injection group as a fixed factor and individual mouse as a random factor; multiple comparisons were accounted for using the sequential Šídák test. Microvessel density analyses were performed using a GLMM with the same random-effects structure. Differences in scRNA-seq cluster composition between tissues were assessed by chi-square test. A p-value < 0.05 was considered statistically significant. Specific sample sizes (n), statistical tests, and p-values are reported in the Figure Legends.

