## Supplementary figures and images for "Generation of functional vasculature from engraftable human pluripotent stem cell-derived progenitors"

### Supplemental Figure 1

#### Supplemental Figure 1.

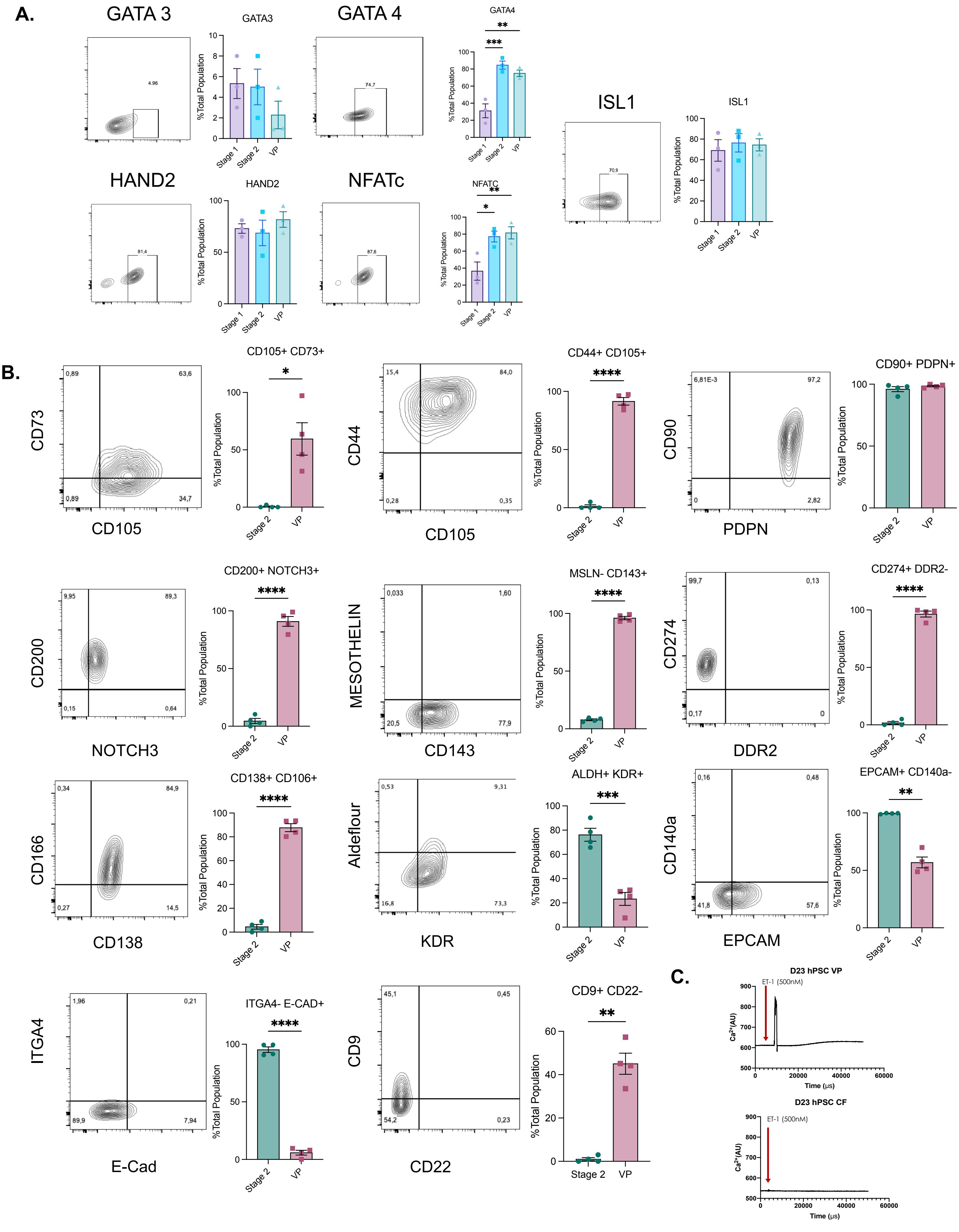

### Supplemental Figure 2

##### Supplemental Figure 2.

**A.**

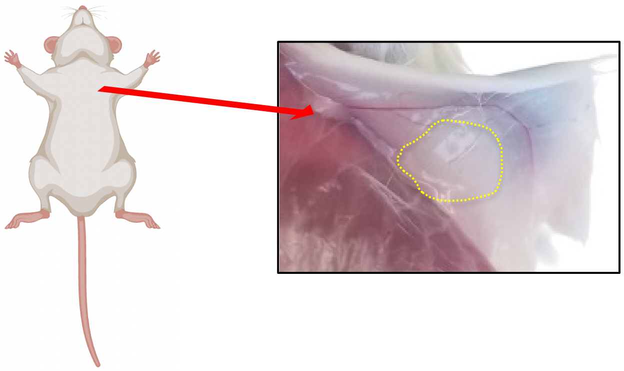

**B.**

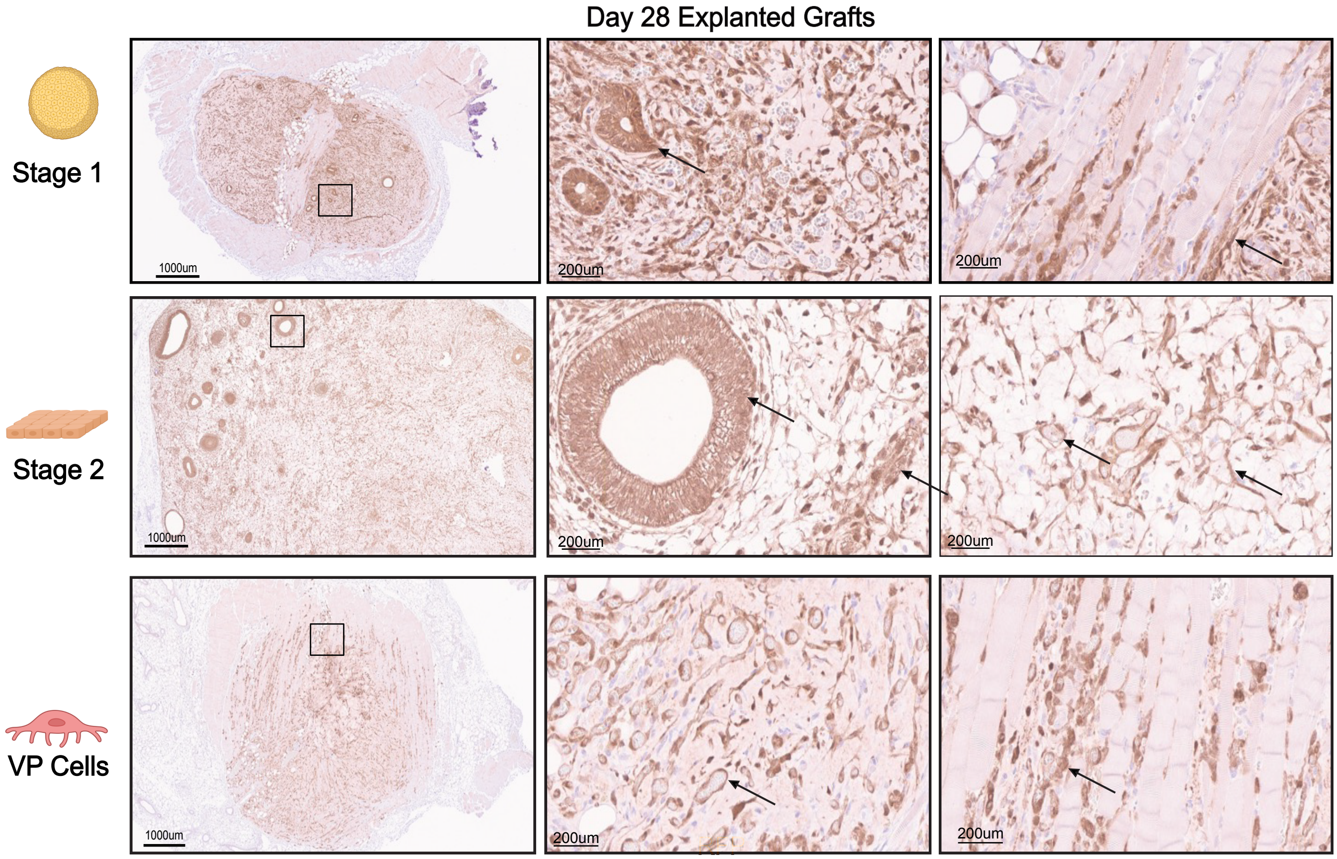

### Supplemental Figure 3

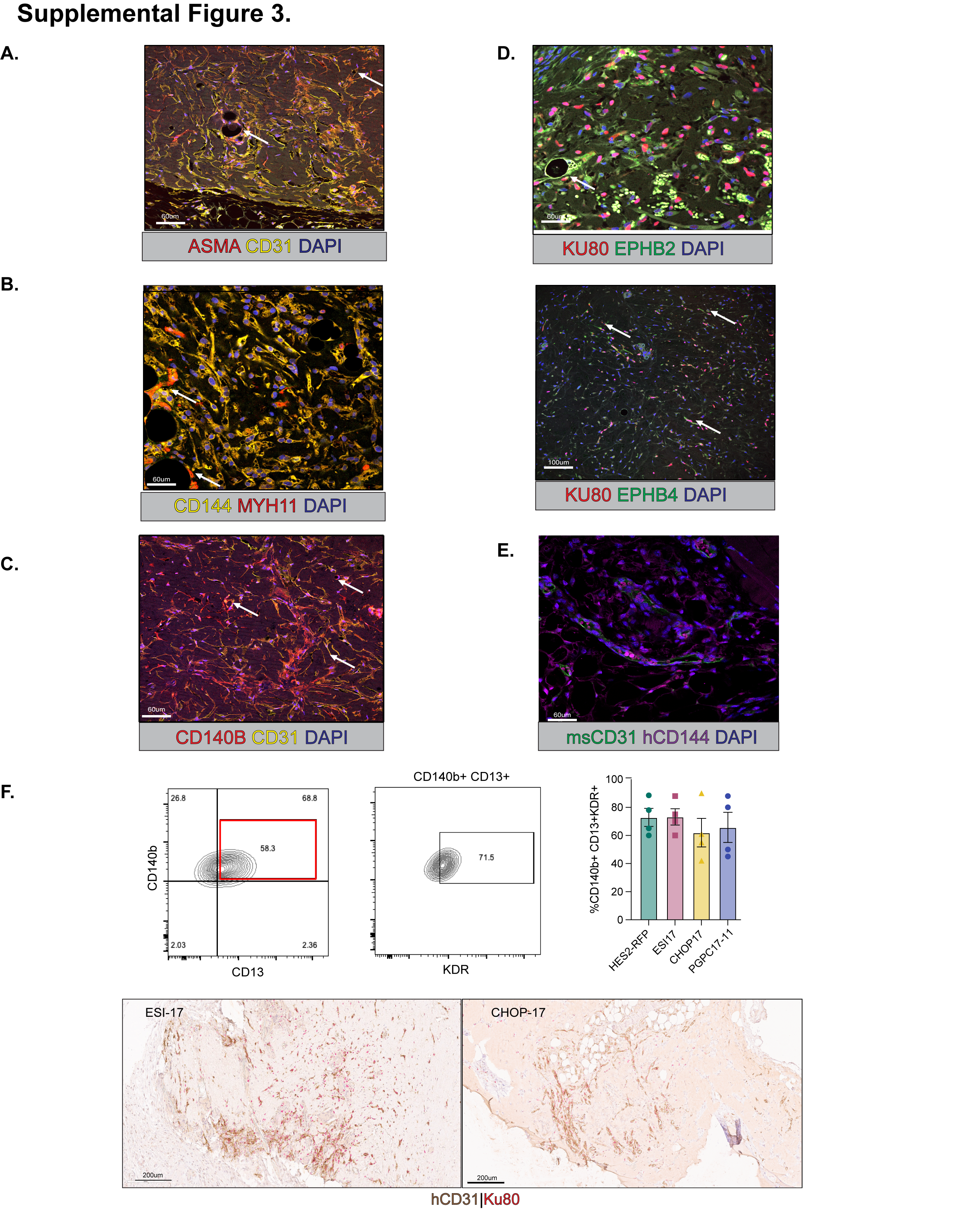

### Supplemental Figure 4

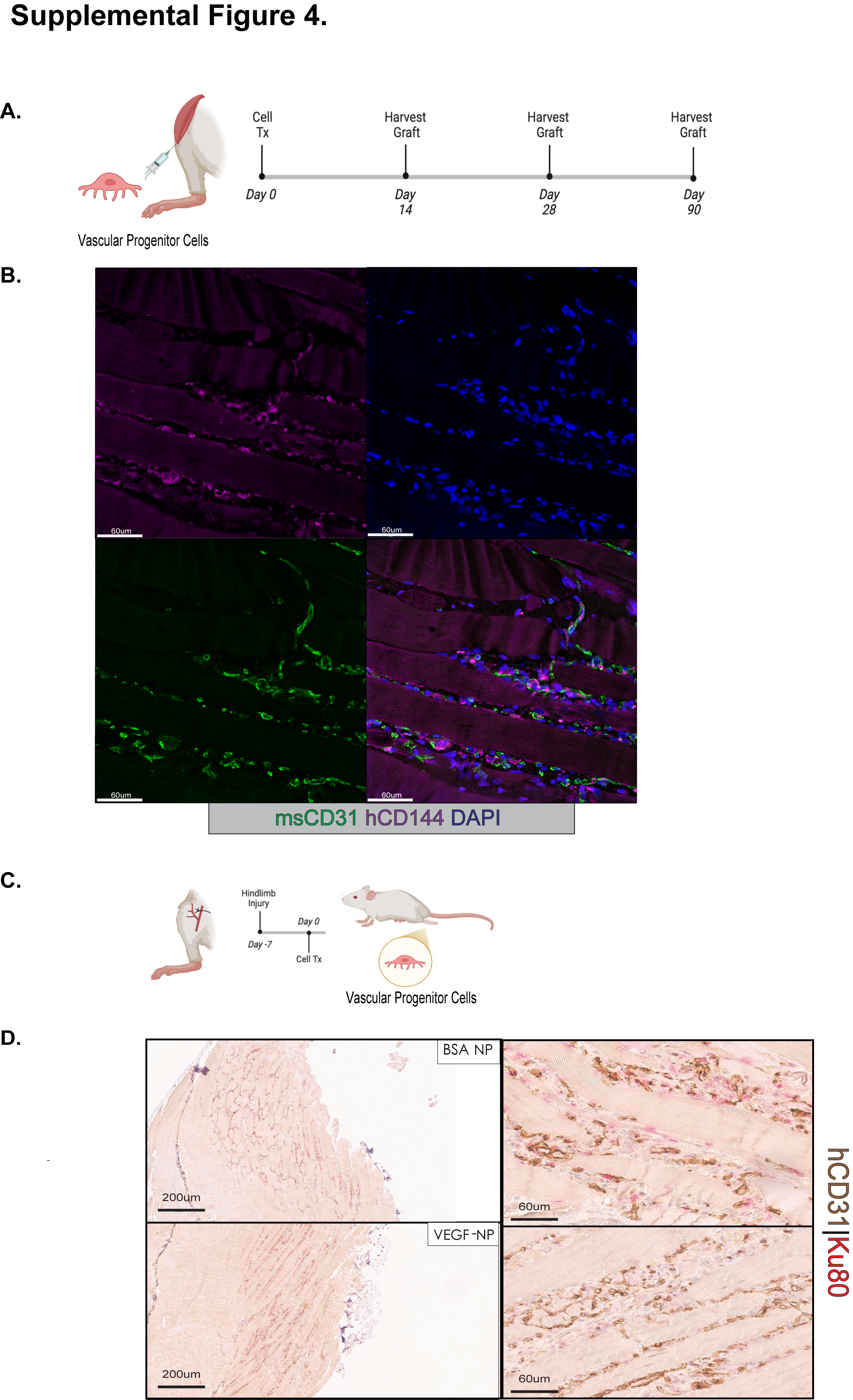

### Supplemental Figure 5

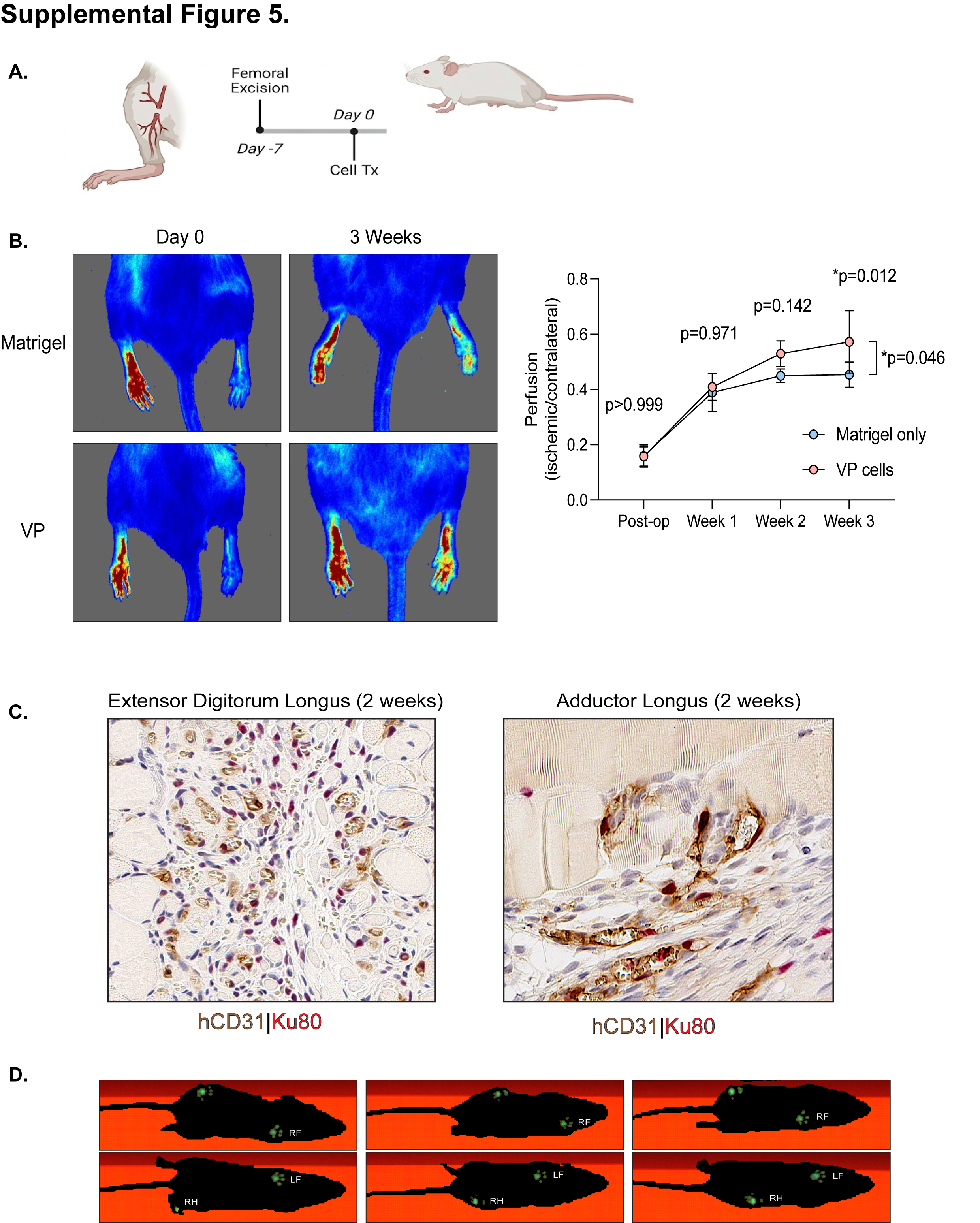

### Supplemental Figure 6

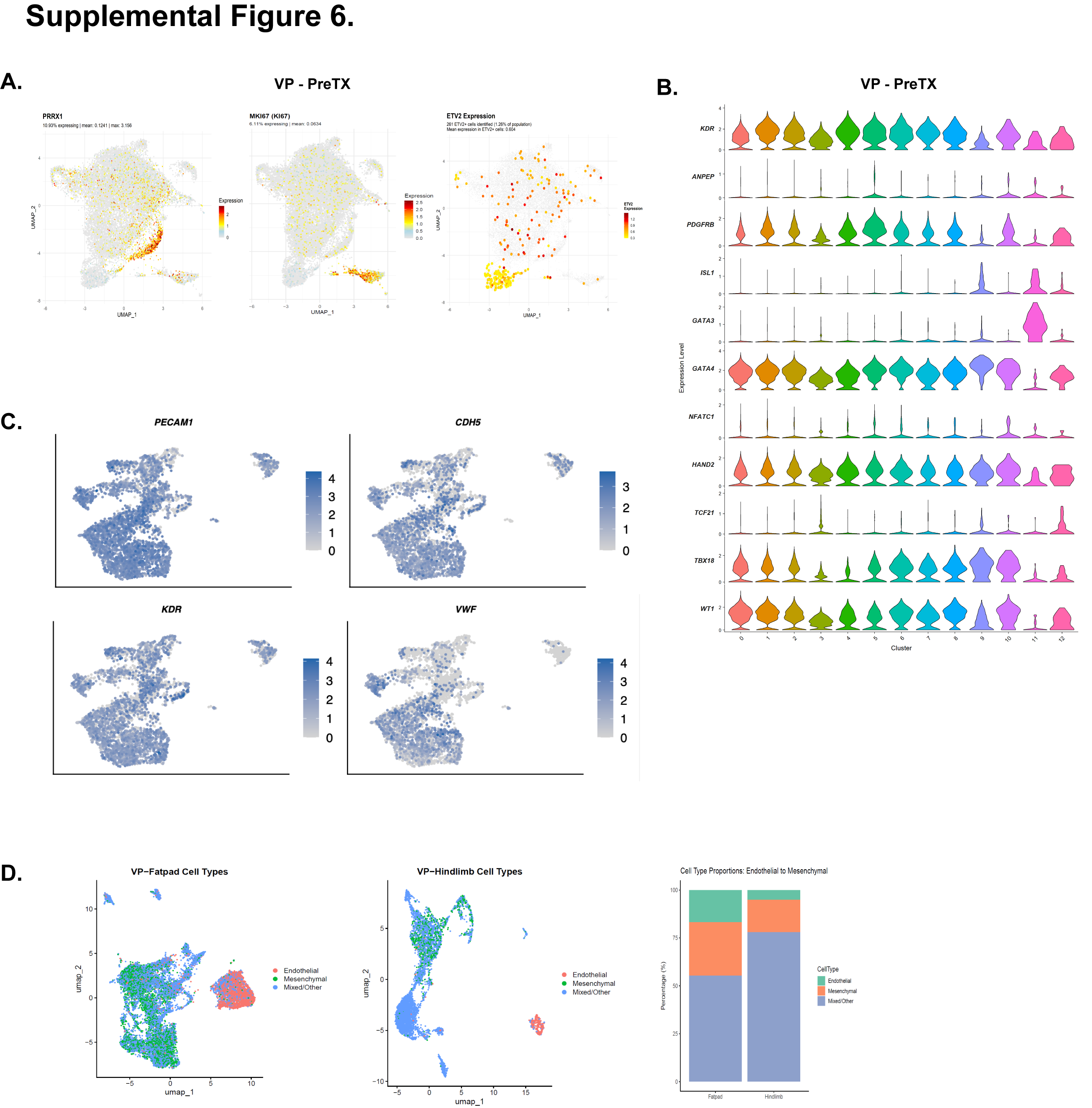

### Supplemental Figure 7

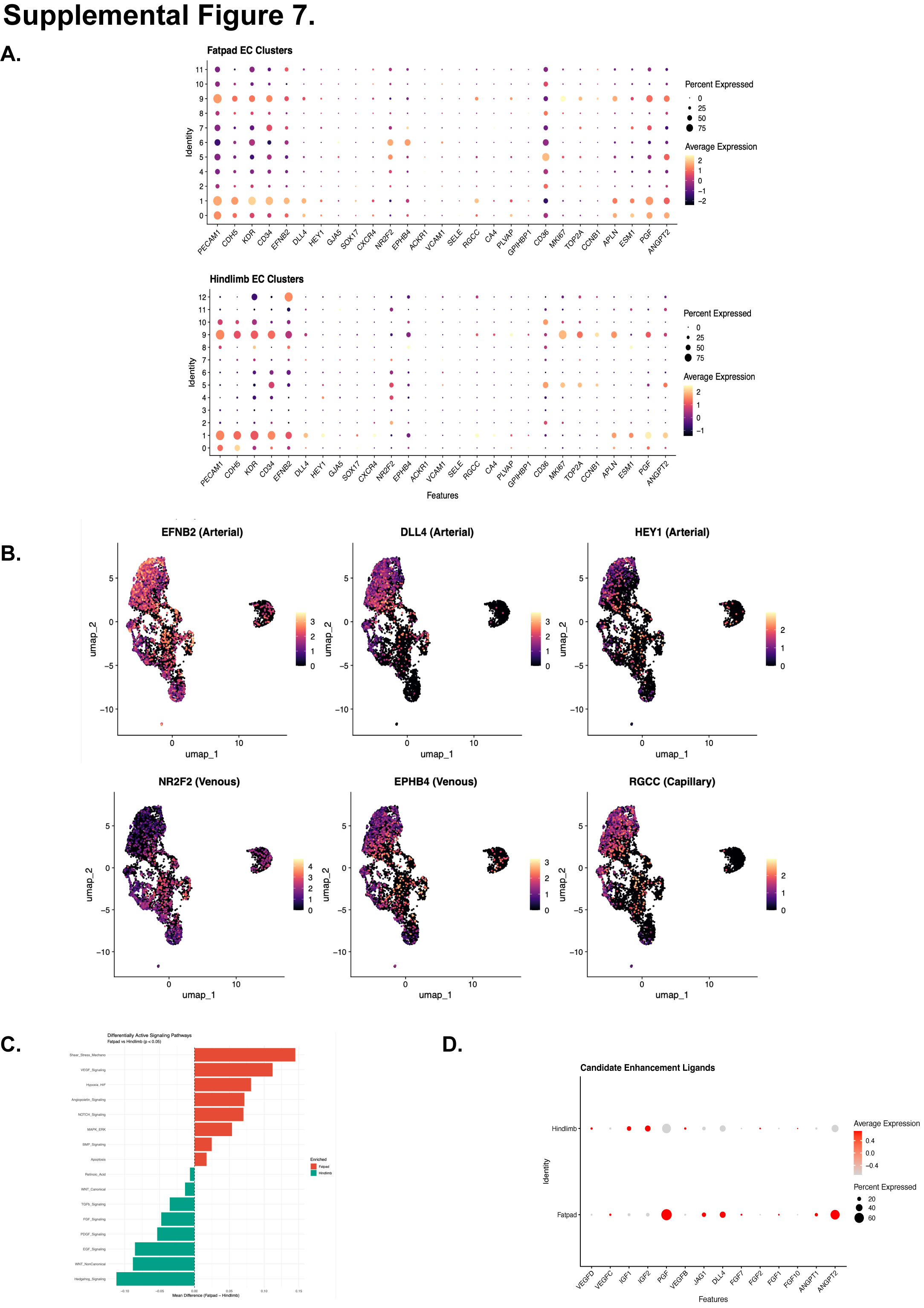
